# Transcriptomes of peripheral blood mononuclear cells from juvenile dermatomyositis patients show elevated inflammation even when clinically inactive

**DOI:** 10.1101/2021.05.07.443007

**Authors:** Elisha D.O. Roberson, Rosana A. Mesa, Gabrielle A. Morgan, Li Cao, Wilfredo Marin, Lauren M. Pachman

## Abstract

In juvenile dermatomyositis (JDM), the most common pediatric inflammatory myopathy, weakness is accompanied by a characteristic rash that often becomes chronic, and is associated with vascular damage. We hoped to understand the molecular underpinnings of JDM, particularly in untreated disease, which would facilitate the identification of novel mechanisms and clinical targets that might disrupt disease progression. We studied the RNA-Seq data from untreated JDM peripheral blood mononuclear cells (PBMCs; n=11), PBMCs from a subset of the same patients when clinically inactive (n=8/11), and separate samples of untreated JDM skin and muscle (n=4 each). All JDM samples were compared to non-inflammatory control tissues. The untreated JDM PBMCs showed a strong signature for type1 interferon response, along with IL-1, IL-10, and NF-κB. Surprisingly, PBMCs from clinically inactive JDM individuals had persistent immune activation that was enriched for IL-1 signaling. JDM skin and muscle both showed evidence for type 1 interferon activation and genes related to antigen presentation, and decreased expression of genes related for cellular respiration. Additionally we found that PBMC gene expression correlates with disease activity scores (DAS; skin, muscle, and total domains) and with nailfold capillary end row loop number (an indicator of microvascular damage). This included *otoferlin*, which was significantly increased in untreated JDM PBMCs and correlated with all 3 DAS domains. Overall, these data demonstrate that PBMC transcriptomes are informative of molecular disruptions in JDM and provide transcriptional evidence of chronic inflammation despite clinical quiescence.

## 1. Introduction

Juvenile dermatomyositis (**JDM**), despite being a rare disease (1), is the most common inflammatory myopathy of childhood (2). It is a systemic, autoimmune process characterized by symmetrical proximal muscle weakness and a typical rash on the face that may cross the nasal bridge, exhibit a shawl-like distribution, or occur on the extremities (3). The rash is often exacerbated on extensor surface skin over joints, such as over knuckles, the front of the knees, and behind the elbows. Classical diagnostic criteria includes elevation of muscle-derived enzymes: creatine phosphokinase (**CPK**), aldolase, lactic acid dehydrogenase (**LDH**), and serum glutamic oxaloacetic transaminase (**SGOT**) (4), which may normalize with a longer duration of untreated disease (5). Myositis Specific Antibodies (**MSA**) are often associated with specific clinical symptoms and outcomes in children with JDM (6). The most common MSA, Tif1- γ (p155/140), present in about 30% of children with JDM, is associated with a chronic relapsing disease course. Disease activity in children with JDM is associated with systemic damage, including microvascular alterations (evaluated by the nail fold capillary end row loop number), and the children often develop premature cardiovascular disease later in life (7, 8). It is unclear whether vascular disease in JDM is driven by the child’s genetic background (9) in addition to duration of untreated disease, overall disease course, and / or the choice of drugs to treat the disease. Some of this ambiguity could be resolved by the development of quantitative molecular biomarkers of disease activity, which could include RNA expression biomarkers. In the current study we first assessed RNA-Seq data from peripheral blood mononuclear cells (**PBMCs**) to identify pathways disrupted in children with JDM prior to treatment when compared to healthy pediatric controls, and then to determine if this signature changed once the child became clinically inactive. We then compared the PBMCs profiles with those obtained from a smaller set of JDM skin and muscle samples to assess the specificity tissue-associated transcriptional signatures and whether more easily obtained PBMCs contain representative signatures of disease activity.

## 2. Results

### 2.1. Sample demographics and clinical information

This study had 2 separate cohorts of children. The first cohort had 3 groups: healthy controls with no known immunological comorbidities (**control**; n=12), children with active JDM prior to treatment (**untreated**; n=11), and later samples from a subset of these children who clinically responded to immunosuppressive treatment (**inactive**; n=8), shown in **Table 1**. All 8 inactive samples were from individuals in the untreated cohort that achieved clinical inactivity. The p155/140 Myositis Specific Antibody (**MSA**) was the most frequent (n=5/11), and 4 children were negative for all tested MSA. The untreated group had average Disease Activity Scores (**DAS**) for skin (**DAS-S**) and muscle (**DAS-M**) each greater than 5.0 (moderately active) and total scores (**DAS-T**) greater than 11.0 (out of a possible 20.0) (10). Inactive samples were selected at a time point when all clinical scores averaged less than 1.0. For the clinically inactive cohort, 4 children were off medical treatment, 1 was tapering a small dose of oral prednisone, and 3 were still undergoing active treatment (combinations of prednisone, mycophenolate mofetil, hydroxychloroquine, and rituxamab; **Table ST2**).

**Table 1.**
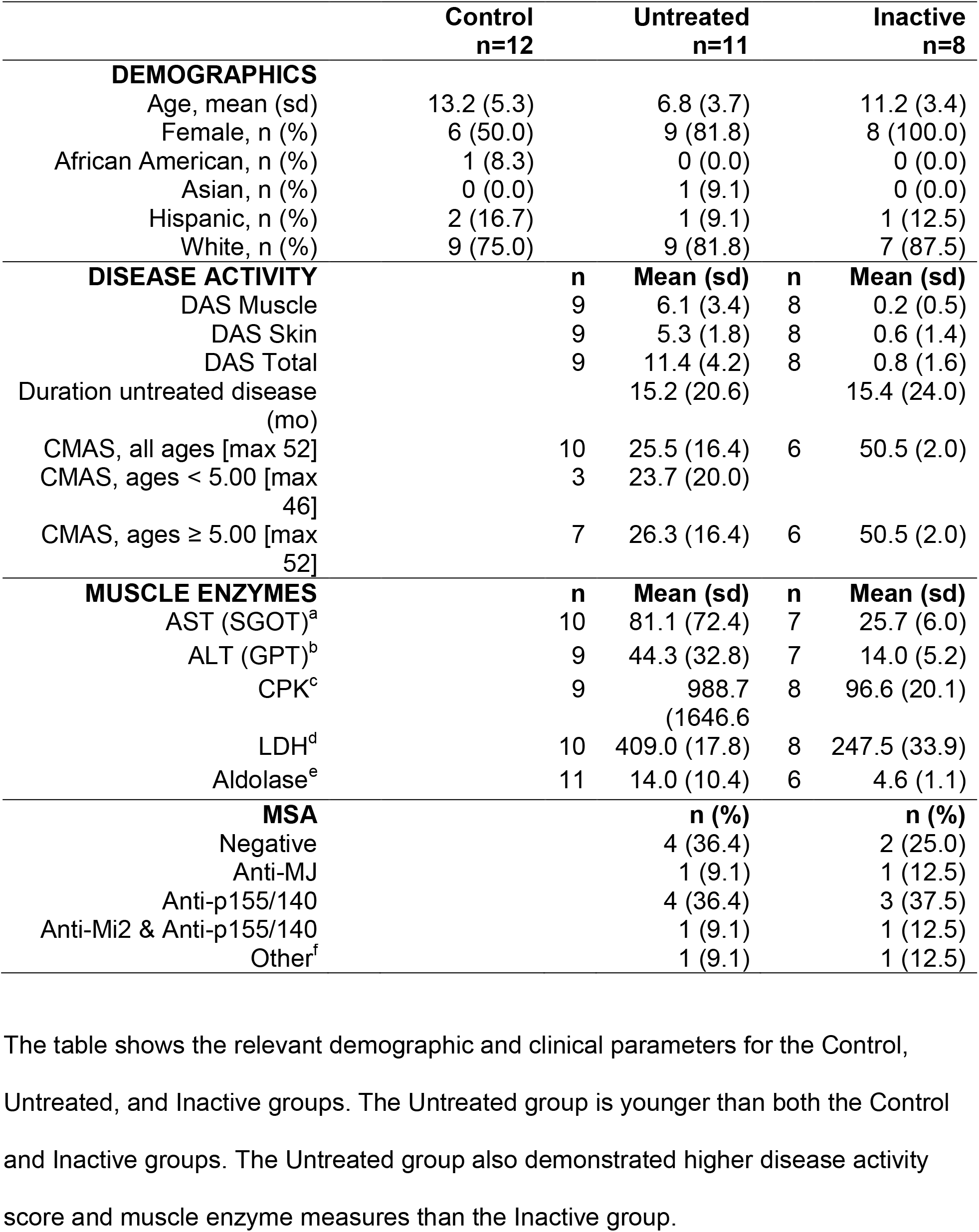
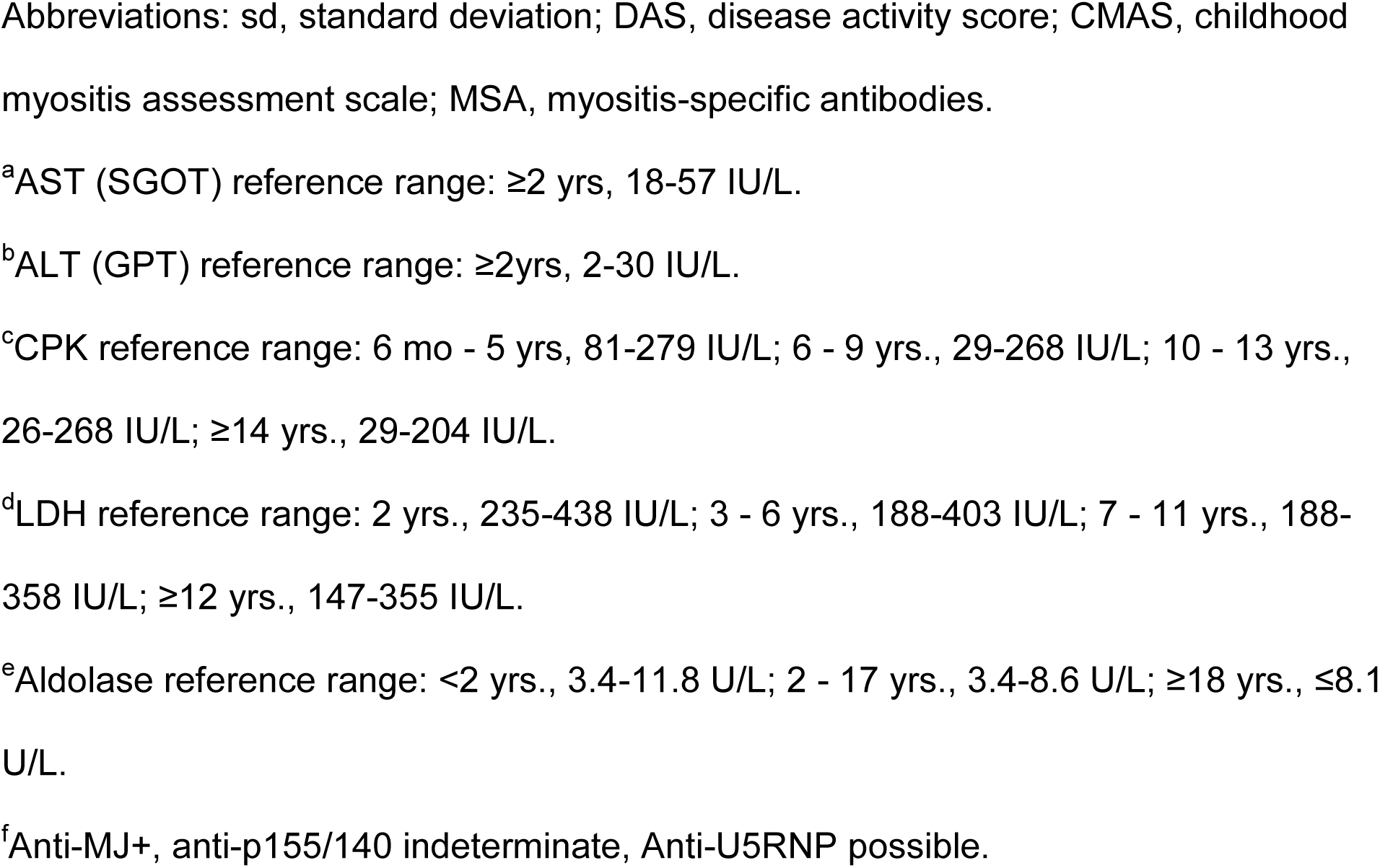
Untreated / Inactive demographics & clinical information.

The second cohort was separate and consisted of skin and muscle samples from children with JDM and otherwise healthy children going to surgery for correction of their idiopathic scoliosis or post-spinal kyphosis. Control skin and muscle samples were not intentionally matched. For each tissue, we had 4 JDM samples (**Table 2**) and 5 controls. The JDM skin and muscle samples were matched. The JDM muscle was obtained at the time of diagnostic muscle biopsy. A sliver of overlying skin was obtained at the same time, but was not selected for having a rash.

**Table 2.**
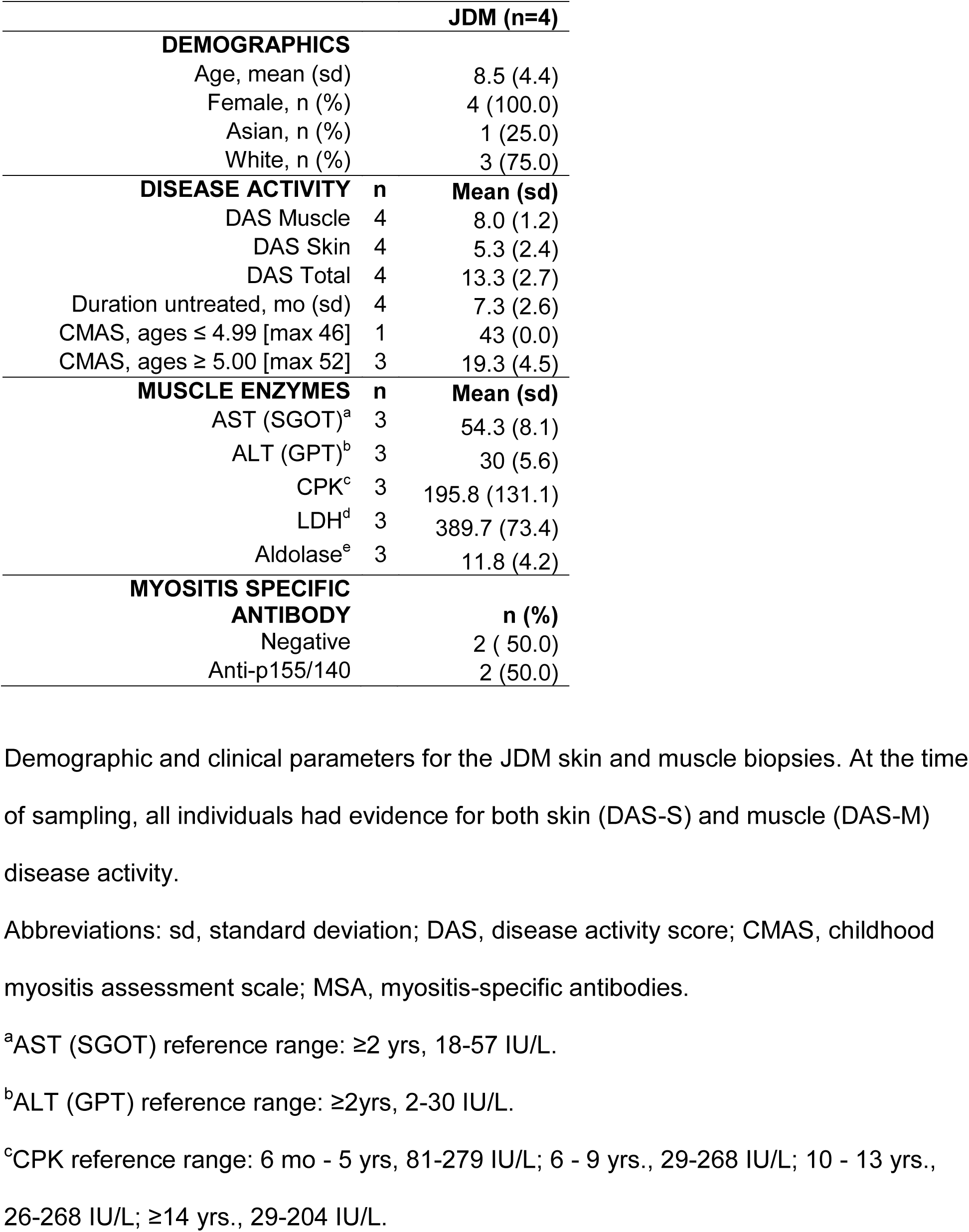
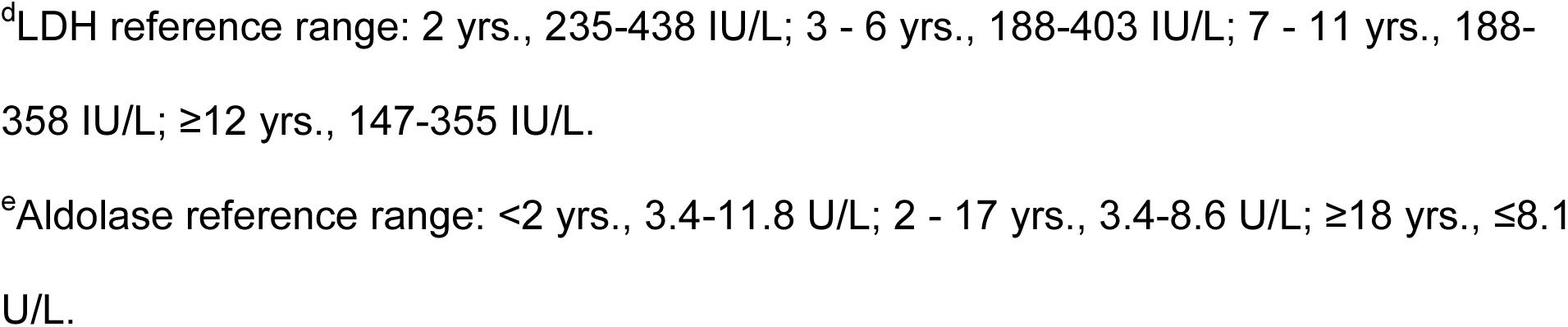
Skin and muscle JDM patient demographics.

### 2.2. Increased type 1 interferon-responsive gene expression characterizes PBMCs from untreated JDM

A total of 301 genes were differentially expressed (**DE**) for the comparison of untreated JDM PBMCs with controls (**Fig. 1A**; **Table ST3**). This included 203 genes increased in untreated (195 at least 1.5 fold) and 98 decreased (93 at least −1.5 fold). Interferon-Induced Protein 44 Like (*IFI44L*) had the most significant increase in untreated JDM with a 10.34 fold-change (**FC**). *IFI44L* is an interferon stimulated gene that is dysregulated in other autoimmune / inflammatory conditions, including multiple sclerosis (11), rheumatoid arthritis (12, 13), and Sjogren’s syndrome (14). Other interferon-related genes had significant increases in untreated JDM as well, including Interferon-Induced Protein with Tetratricopeptide Repeats 2 (*IFIT2*; 7.92 FC), 2’-5’ Oligoadenylate Synthetase-Like (*OASL*; 6.90 FC), and Interferon-Induced Protein 44 (*IFI44*; 4.89 FC). There was also a marked increase (3^rd^ most significant) in otoferlin (*OTOF*; 23.60 FC). The increase of *OTOF* in the untreated JDM PBMCs compared to controls was confirmed by RT-qPCR (**Fig. 2**; FC 11.02 ± 5.12; P = 5.6 x 10^-4^).The ferlin families of proteins are membrane anchored, aid in membrane healing, and are involved in vesicle fusion (15). Otoferlin is known to facilitate calcium flux in inner ear hair cells and has lipid binding properties (16–18). The gene *EIF2AK2*, also called PKR (RNA-dependent protein kinase R), was increased 2.74-fold. Published studies document that the PKR protein interacts with several parts of the inflammasome, and is critical for its activation (19).These top DE genes suggested that the primary signal was from type 1 interferon response. This was confirmed by taking the genes with significantly increased expression and testing for pathway enrichment using gProfileR (20). The top enriched pathways for genes with increased expression included type 1 interferon response, interferon signaling, and response to virus (**Fig. 1B**; **Table ST4**). The increased expression genes were also enriched for ISGF-3 transcription factor sites (TF:M00258_1) in their proximal promoters (n=8 genes; adj. p-value 2.59 x 10^-7^), along with signals for IRF1-5, IRF7-9, and STAT2. Consistent with an up-regulated immune response, there were enrichments for the migration and chemotaxis of lymphocytes (GO:0048247), monocytes (GO:0002548), neutrophils (GO:0030593), and natural killer cells (GO:0035747). Cytokine and chemokine enrichments (GO:0042379, GO:0008009, GO:0005125) were driven primarily by increases in *CCL2-4*, *CCL3L1*, and *CCL4L2*, along with CXCL genes *CXCL1-3*. Of the differentially expressed cytokines, terms for IL1 signaling (GO:0071347, GO:0070555) and IL1 receptor binding (GO:0005149) were enriched, mostly due to overexpression of *IL1A*, *IL1B*, *IL1RN*, and *IL6*. Increased expression of *NFKB1* and *NFKBIA* led to an enrichment of the NFKB (KEGG:04064) and TNF signaling pathways (KEGG:04668).

**Figure 1.**
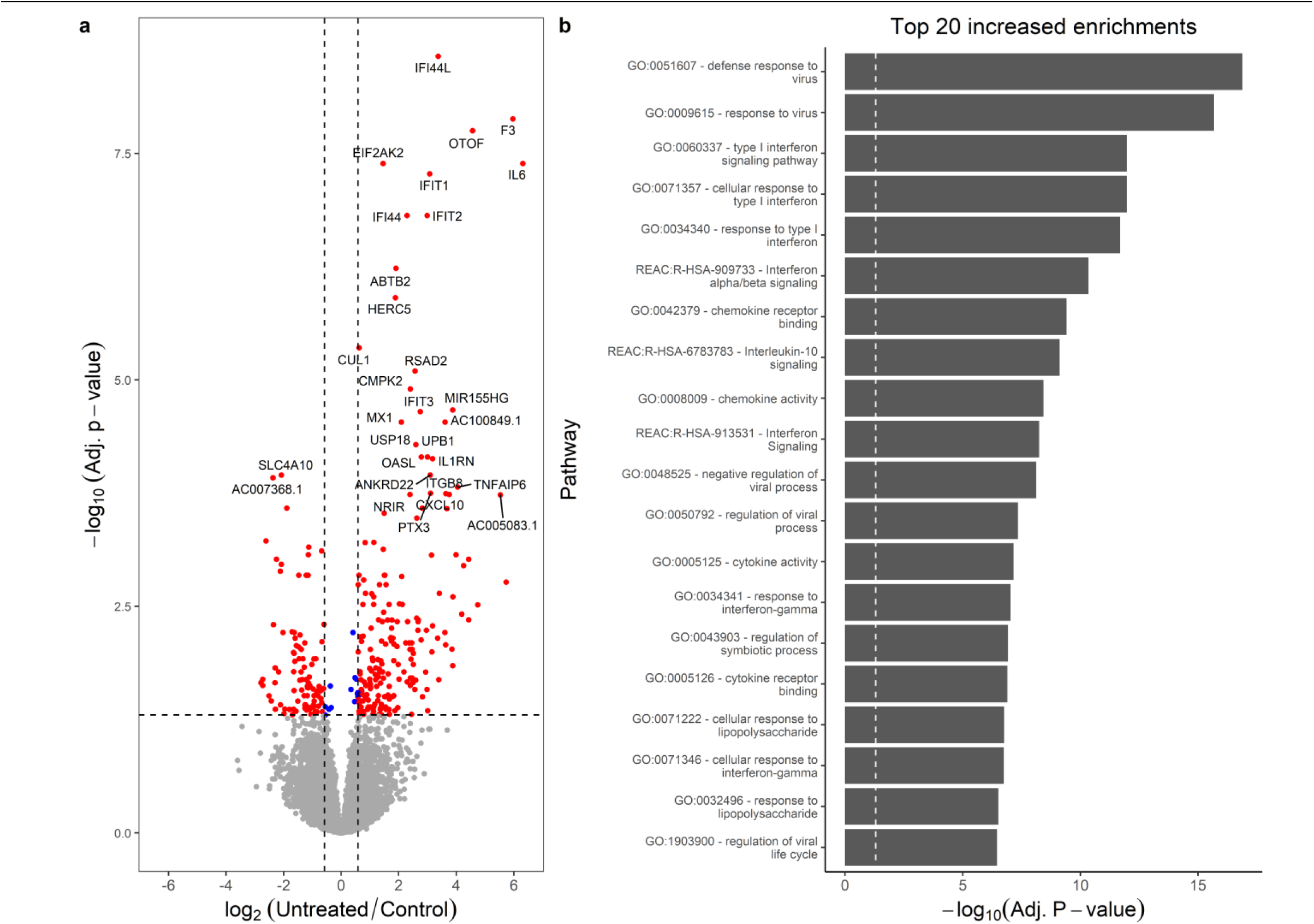
Untreated JDM has enrichment of type 1 interferon signaling. **a.** Volcano plot for samples from untreated JDM patients versus controls. The x-axis shows the log_2_ fold-change, and the y-axis shows the false-discovery rate corrected p-value. The two vertical lines indicate the cutoff points for a 1.5-fold change. The horizontal line indicates the 0.05 adjusted p-value cutoff. Some of the most significant genes are individually labeled. Insignificant points are colored gray. Significant points are labeled red or blue depending on whether the fold-change met a 1.5-fold threshold. There are some significantly decreased genes, but by far the most significant changes are increases in inflammatory genes, specifically interferon signaling. **b.** Significantly enriched pathways for genes with increased in untreated JDM from gProfileR. There are enrichments interferon signaling, cytokines, chemokines, and IL-10.

**Figure 2.**
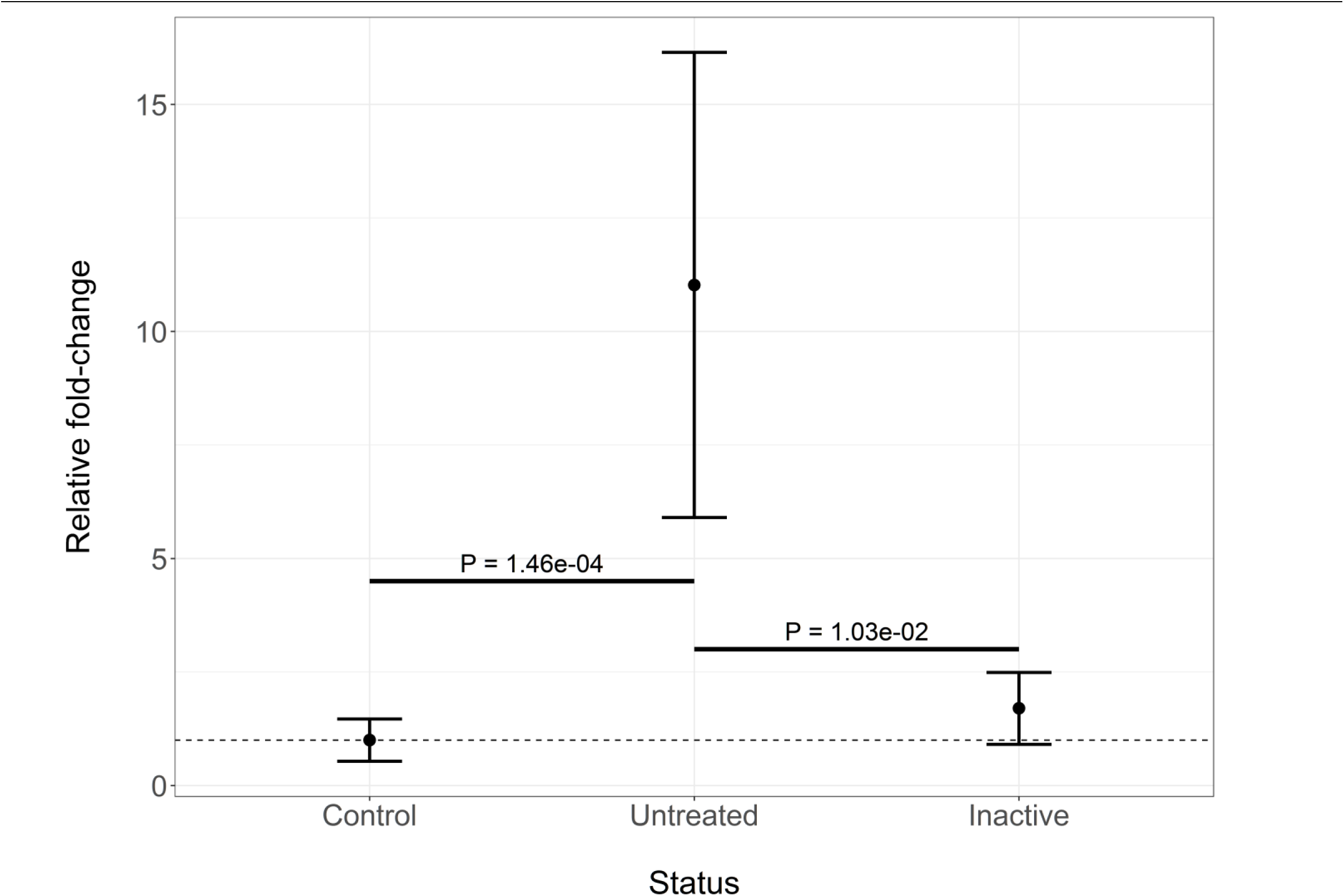
Otoferlin is increased in untreated JDM PBMCs. RT-qPCR confirmation of increased expression of *otoferlin* in JDM. The x-axis shows the disease status of the samples, which includes control (n=13), untreated JDM (n=11), or inactive JDM (n=7). The y-axis shows the mean and standard deviation of gene expression relative to controls. The untreated samples have significantly increased *otoferlin* expression compared to both control and inactive PBMCs.

There were fewer genes with significantly decreased expression, and they did not fall consistently into known pathways (**Table ST5**). There were only two significant enrichments in Gene Ontology (**GO**) cellular compartments (**CC**), and only one was driven by decreases in several genes: cell cortex (GO:0005938). The lack of consistent pathway enrichments makes the decreased genes more difficult to interpret and requires some examination of individual genes. Individual genes with decreased expression in untreated JDM included Solute Carrier Family 4 Member 10 (*SLC4A10*; −4.53 FC), ST8 Alpha-N-Acetyl-Neuraminide Alpha-2,8-Sialyltransferase 1 (*ST8SIA*, −5.04 FC), and Acrosin Binding Protein (*ACRBP*; −1.94). Of importance, knockdown of *ST8SIA* interferes with the induction of autophagy in the 2FTGH human fibroblast cell line. *ACRBP* is decreased in Ly6c^low^ monocytes after treatment with rosiglitazone, a PPARgamma agonist (21). *IL23R* was reduced as well (−4.76 FC). A heterodimer of IL23R and IL12RB1 is required for canonical IL23A signaling. *IL12RB1* was not reduced significantly (0.58 adjusted P-value). ADP-ribosylation factor-like3 (*ARL3*) had a −1.52 fold reduction in untreated JDM. Overexpression of *ARL3* in the human HEK293T cell line is associated with a significant increase in markers associated autophagy (22), so decreased expression might be associated with decreased autophagy. The finding of multiple genes related to autophagy having decreased expression raises the interesting question of whether disrupted autophagy is an important mediator of JDM onset and progression.

### 2.3. Clinically inactive JDM patients have persistent immune activation with enrichment for genes associated with IL-1, IL-10, and NF-κB signaling

The individuals classified as inactive (n=8 females) had markedly reduced clinical disease activity compared to their untreated sample time point. We hypothesized that this clinical response would result in their transcriptomes reverting to a control-like state. For this test we determined the differences as untreated / inactive. If a gene’s fold-change was in the same direction as the untreated / control comparison, the signature must have improved with treatment. If the fold-change was in the opposite direction, then the expression level for that gene had worsened. Genes that were DE in the untreated / control comparison, but unchanged in the untreated / inactive comparison were still dysregulated in the inactive sample. The paired test between the untreated samples and the matching inactive sample with DESeq2 (Expression ∼ Individual + Status) showed there were 217 significantly different genes. This consisted of 91 genes with increased (90 at least 1.5-fold) and 126 decreased (123 at least −1.5-fold) expression in the untreated sample compared to clinical inactivity (**Fig. 3**; **Table ST6)**.

**Figure 3.**
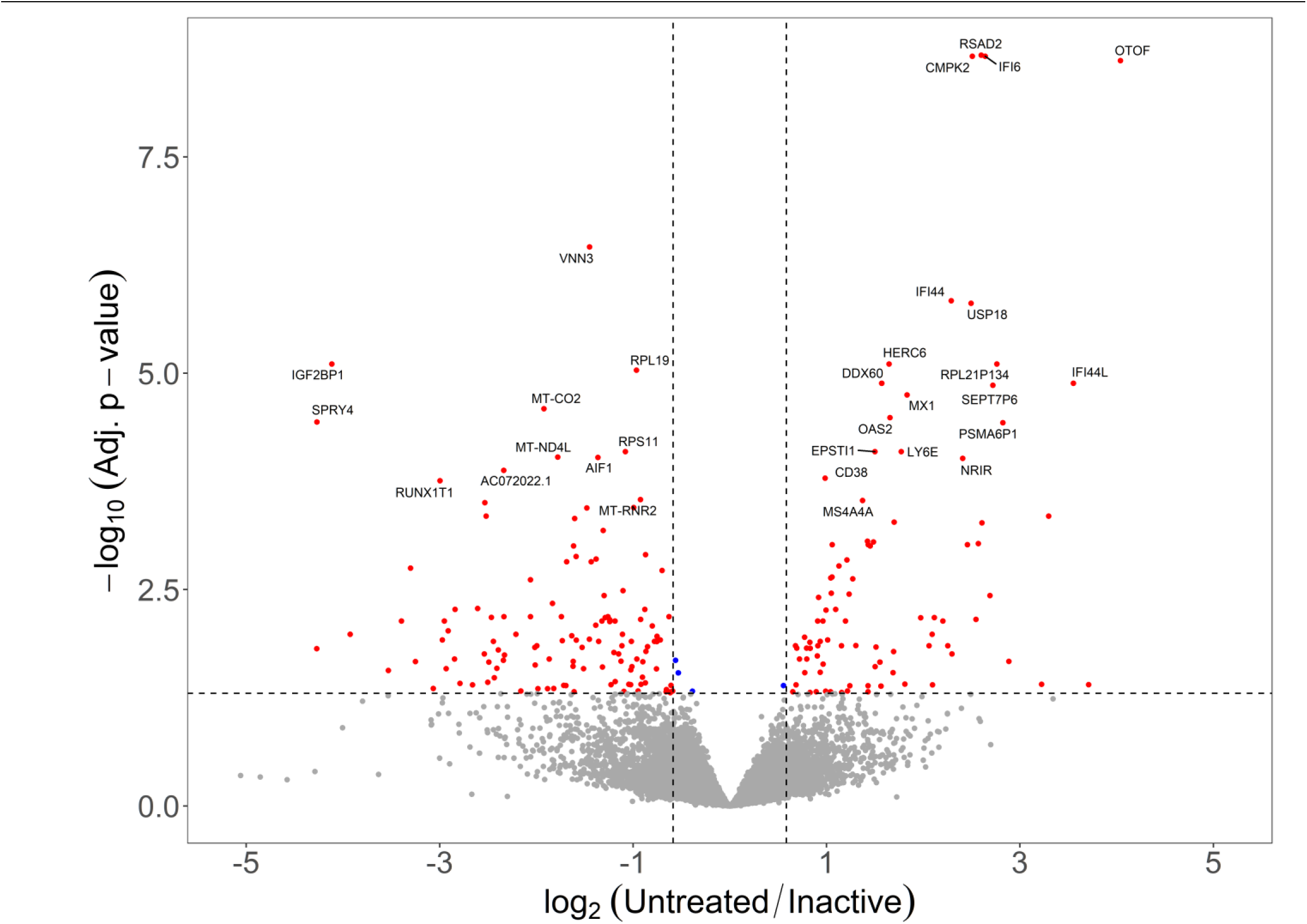
Inactive JDM PMBC samples do not revert to control-like gene expression. Volcano plot with log_2_ fold-change on the x-axis and –log_10_ of the false-discovery rate corrected p-value. The vertical lines indicate a minimum 1.5-fold change. The fold-change is represented as untreated / inactive. Genes that are in the upper right area were increased in untreated, but resolved with inactivity. Genes in the upper left area were decreased in untreated JDM and resolved with clinical inactivity. Most genes fall below the horizontal significance cutoff line (grey dots), indicating the untreated and inactive PBMCs are not different from each other for the gene.

Some of the top DE genes from the untreated vs. control comparison, such as *RSAD2*, *CMPK2*, *IFI6*, *OTOF*, *IFI44*, and *MX1*, did resolve when the individual entered clinical inactivity (**Fig. 4**). It’s possible that some of the resolved genes are key drivers of overt clinical progression. Strikingly, the majority of the genes (∼85%) that were differentially expressed when untreated remained altered after the same individual was clinically inactive (**Table 3**; **Fig. 5**). The DE genes in this comparison therefore contain genes that resolved with treatment, as well as genes that were not DE in the untreated vs. control comparison, but are DE in this untreated vs. inactive comparison.

**Figure 4.**
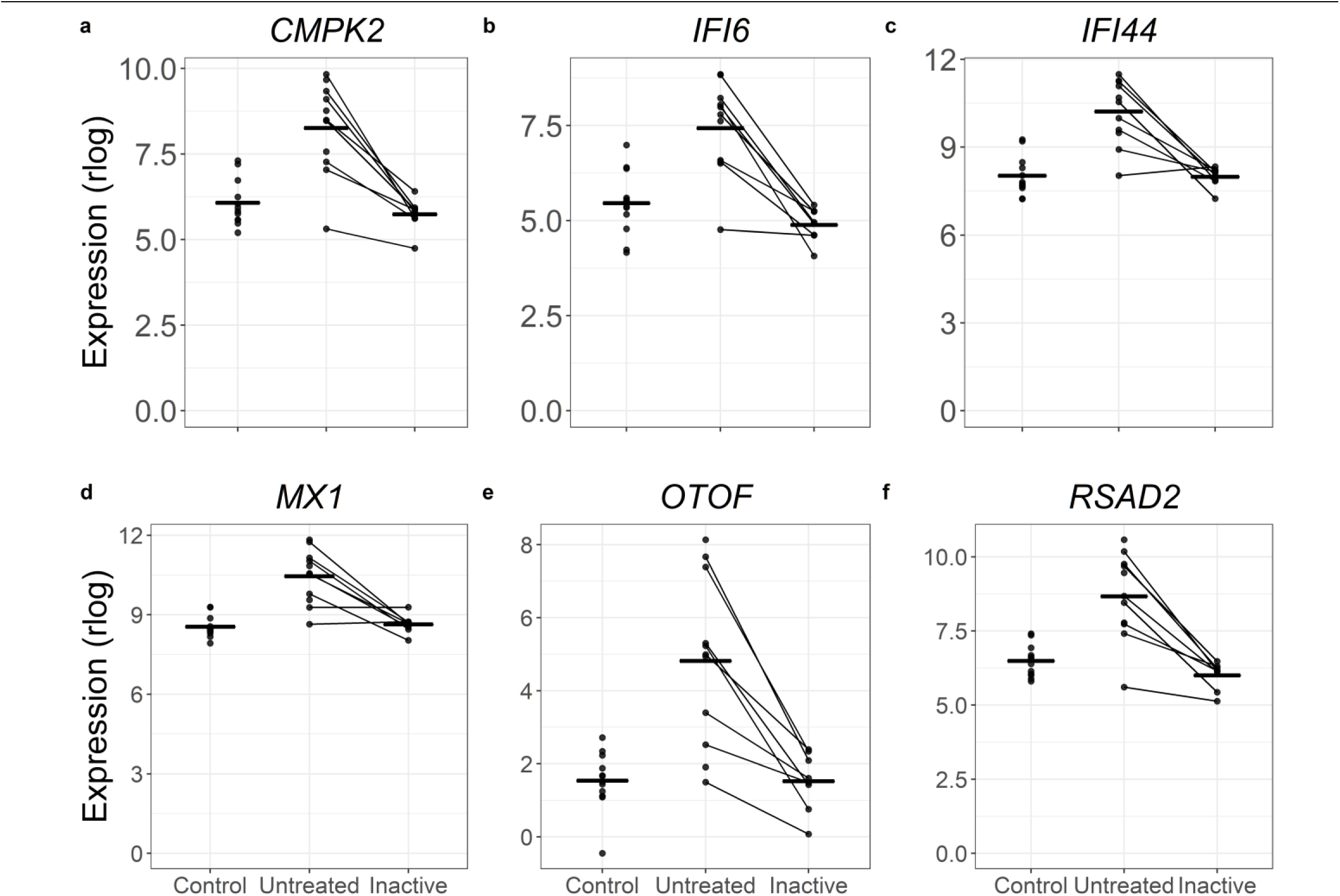
A subset genes increased in untreated JDM resolve when patients are clinically inactive. In each panel the layout is the same. The x-axis shows the status for each sample. The y-axis is the normalized expression (regularized logarithm method) from the RNA-Seq data. Each point is an individual sample. The lines connecting dots indicate sample pairings from the same individual. The horizontal bars are category means. For all these particular genes, the untreated sample PBMCs show substantially increased (and variable) increased expression. When inactive, the expression levels decreased toward the control expression level.

**Figure 5.**
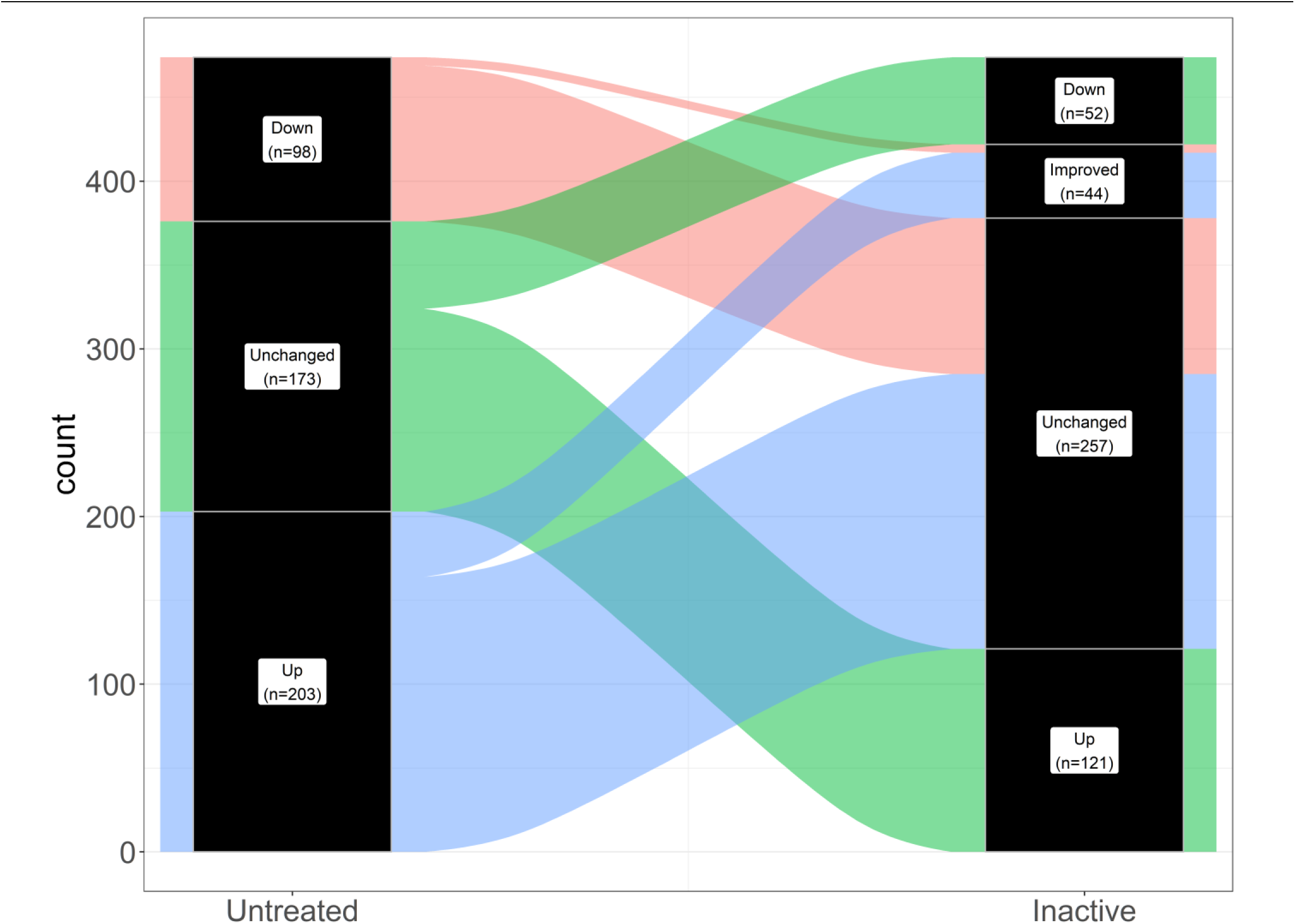
Alluvial plot demonstrating that most genes differentially expressed in untreated samples remain dysregulated in inactivity. The left column (labeled Untreated) represents the status of gene expression for the untreated vs. control comparison. Genes increased untreated samples are filled in blue. Genes decreased in untreated samples vs. controls are filled in light red. Genes unchanged in untreated vs. control, but altered in untreated vs. inactive are filled in green. The right column (labeled Inactive) shows the response of paired samples when they reached clinical inactivity. Out of all the differentially expressed genes in untreated vs. control, only 44 resolved toward control-like expression. Most genes remained altered in the inactive samples.

**Table 3.** Transcriptional response to clinical inactivity.

| **Untreated vs. Control** | **Untreated vs. Inactive** |
| --- | --- |
| Increased (n=203) | Improved (n=39)<br>Unchanged (n=164) |
| Decreased (n=98) | Improved (n=5)<br>Unchanged (n=93) |
| Not DE (n=173) | Increased (n=121)<br>Decreased (n=52) |

Data are shown for two different contrasts: untreated JDM PBMCs vs. controls, and untreated JDM versus inactive JDM. In the untreated JDM versus control comparison, there are three possibilities: 1) increased in untreated JDM, 2) decreased in untreated JDM, or 3) not differentially expressed. We assessed each of these categories to determine whether they changed with inactive disease. This is shown in the second column. Some genes improved toward a control-like state when inactive. Another set of genes that were not differentially expressed in the untreated vs. control comparison were changed when inactive. Overall, most genes that were either increased or decreased when untreated did not improve when patients were inactive.
Abbreviations: DE: differentially expressed; JDM, juvenile dermatomyositis; PBMCs, peripheral blood mononuclear cells.

Additionally, some genes that were not significant in untreated vs. control were significant in untreated vs. inactive. This could be a change in gene expression due to duration of disease or response to specific treatments. It’s worth noting that the power of a paired test can often be greater than a group-wise test since it focuses on changes between conditions rather than pure difference in means. Many of these newly DE genes in the untreated vs. inactive comparison may represent changes that were underpowered for the group-wise test, but well-powered for the paired comparison of untreated vs. inactive. Because these possibilities are more complex than the results of a standard two-sample test, we tested for enrichments separately for improved genes, still dysregulated genes, and newly differentially expressed genes.

For genes that were increased in untreated samples, but improved when they reached clinical inactivity there was a clear signal for viral defense and type 1 interferon (**Table ST7**; GO:0051607, GO:0009615, REAC:R-HSA-909733, GO:0060337, etc).

There were also enrichments for targets of STAT2 and several IRF transcription factors. This suggests that at least some of the interferon activation was suppressed by treatment. For genes that were increased in untreated samples, but still dysregulated when clinically inactive, there were different enriched pathways (**Table ST8**). There were minor themes for type 1 interferon signaling, but the major signals were for IL-10 signaling (REAC:R-HSA-6783783), response to IL-1 (GO:0071347, GO:0070555,

GO:0070498), and NF-KB signaling (KEGG:04064, GO:0007249). This is a particularly important point, since it suggests that different immune axes drive different phases of disease progression. It also suggests that standard physical examination may not be sensitive to persistent inflammation. There were no significant enrichments for genes that were decreased in untreated samples that resolved with clinical inactivity. Similarly, there were only 12 enrichments for genes decreased in untreated JDM that were still decreased with clinical inactivity (**Table ST9**). These pathways were not as informative, primarily composed of pathways related to microtubule transport (GO:0098840, GO:0099118, GO:0010970, GO:0099111) and EGFR signaling (KEGG:01521).

The interpretation of genes that were not differentially expressed in untreated samples, but were significantly different when the JDM was clinically inactive is more difficult. We first tested for enrichments for genes that were increased in inactive samples compared to the untreated samples (**Table ST10**). The enrichments were clearly related to mitochondrial function, including cellular respiration (GO:0098803, GO:0070469, HP:0200125) and oxidative phosphorylation (GO:0006119, WP:WP111, KEGG:00190). Both of these themes were driven by mostly by differential expression of several mitochondrial encoded genes, including *MT-ATP6*, *MT-ATP8*, *MT-CO2*, *MT-CO3*, and *MT-ND1-5*. In this case, it appears that instead of newly increased expression in the inactive sample, the expression was insignificantly decreased in untreated samples and normalized in the clinically inactive samples (**Fig. 6**), supporting the idea that in this case these genes were detected because of the increased power of a paired test.

**Figure 6.**
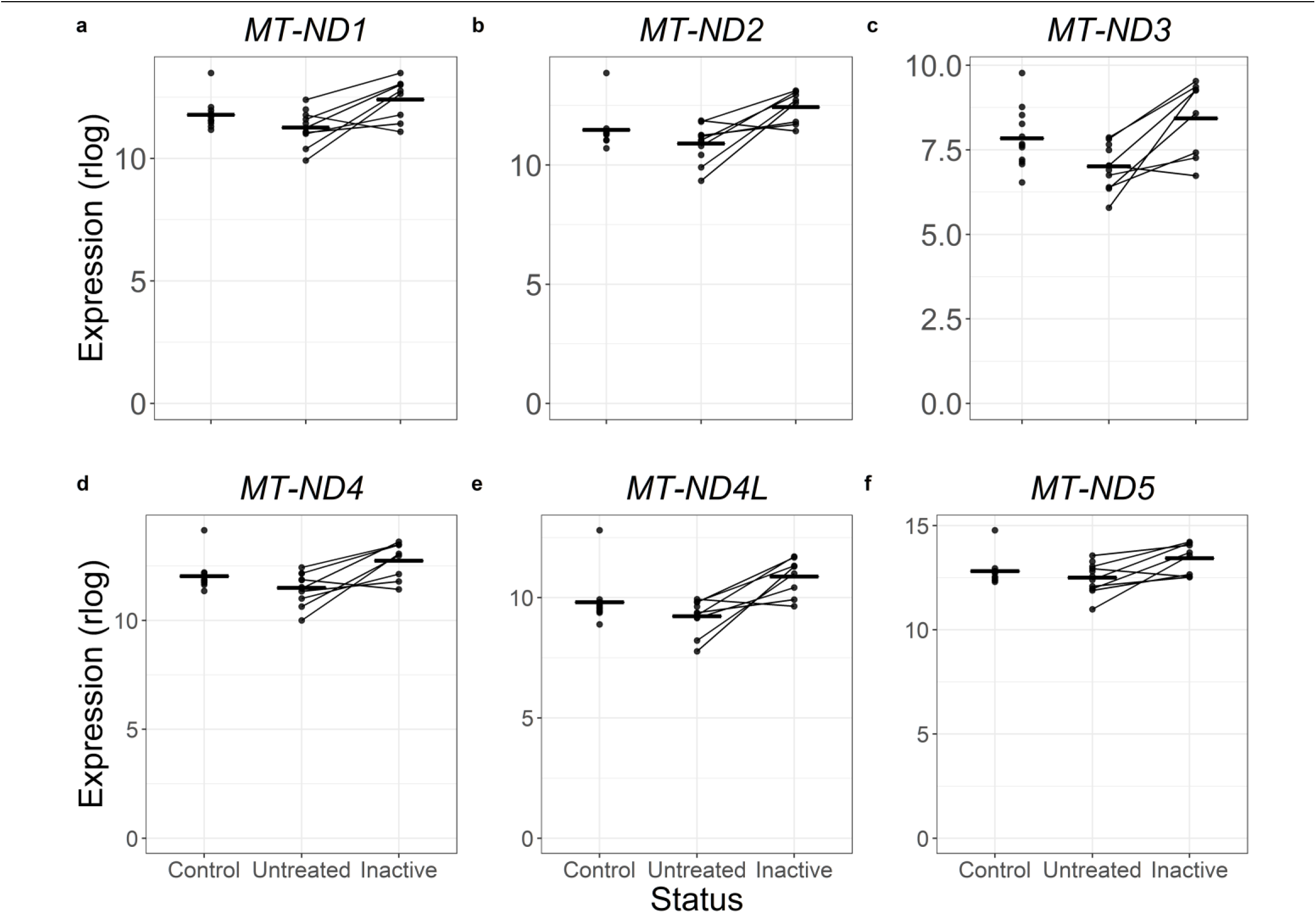
Mitochondrial genes trend toward being decreased in untreated JDM PBMCs, but improve when patients are clinically inactive. Six panels with the same plot arrangement. In each panel, the x-axis shows the disease status of the sample, and the y-axis shows normalized expression via the DESeq2 regularized logarithm. Each dot is a single sample, with lines connecting paired samples. The horizontal bars are group means. The mitochondrial decrease was not a strong signal in the untreated vs. control comparison, but was a major theme in untreated vs. inactive comparison. These plots are able to show a trend toward decreased expression in untreated JDM PBMCs, through the significance was not high. When clinically inactive, the mitochondrial gene expression returned to the control expression range.

Another set of genes was not statistically different in the untreated vs. control comparison, but was decreased in inactive samples when compared to their own untreated samples (**Table ST11**). There were few enriched pathways, but there was a consistent theme of interferon responses. This also appeared to be a case of increased power. Three of the genes driving these themes were *IFI16*, *MX2*, and *OAS1*. Looking across the different status categories, it’s apparent that all three trended toward being increased in the active untreated samples without reaching significance, and that the inactive samples regressed toward control-like expression levels (**Fig. 7**).

**Figure 7.**
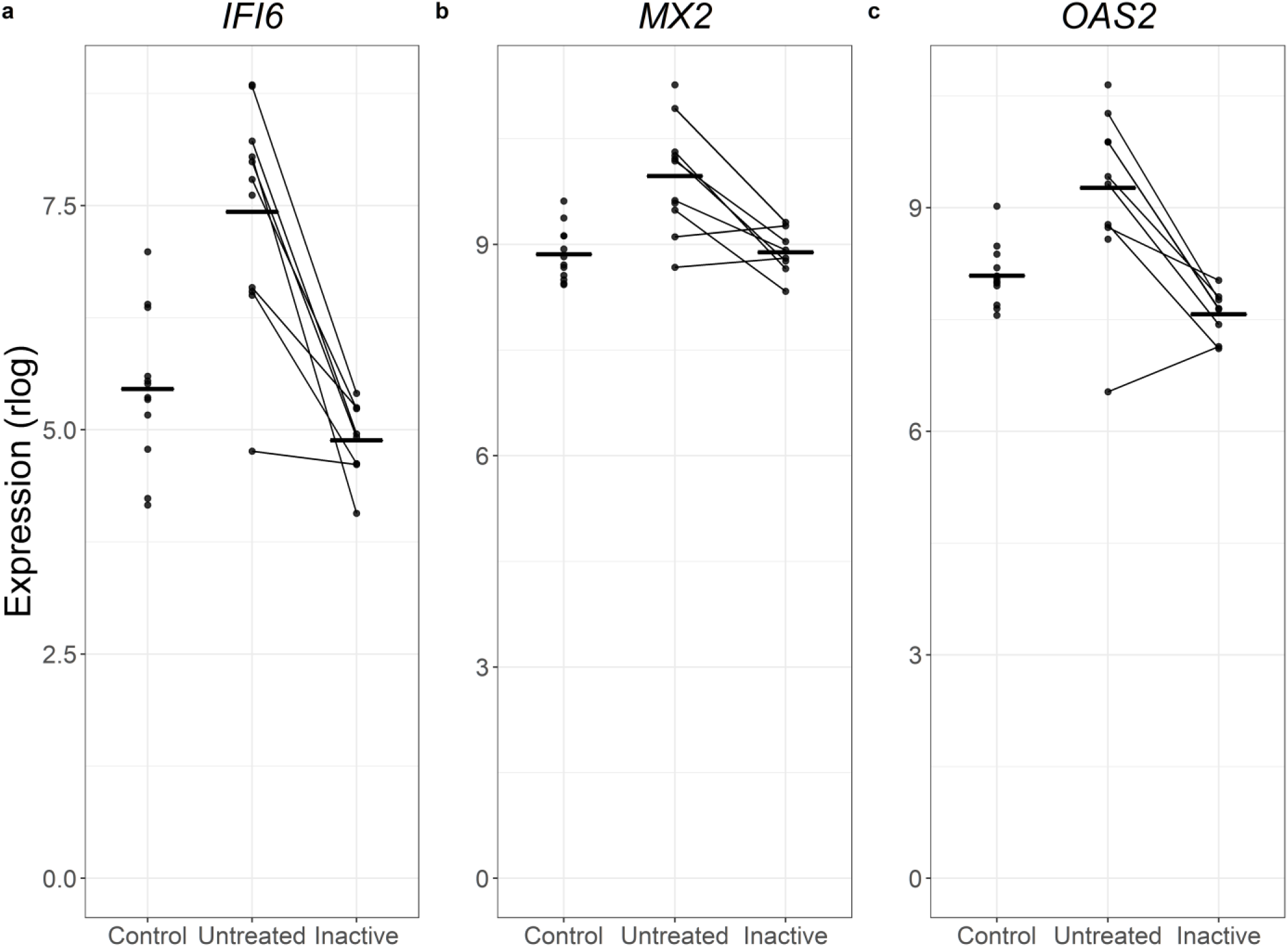
Additional altered interferon-related genes can be identified when comparing untreated JDM PBMCs to inactive PBMCs. Three type 1 interferon signaling-related genes shown in 3 individual panels. In each panel, the status is on the x-axis, the normalized expression (regularized logarithm) is on the y-axis, and the horizontal bars show group means. Each dot is a sample, and lines connect paired samples. These are genes that were not differentially expressed in untreated vs. control, but were differentially expressed in untreated vs. inactive. The pattern suggests a non-significant increase in untreated JDM that resolves with inactivity. This is likely due to greater power for a paired differential expression test.

The group-wise tests are helpful in pointing out shared trends of changes in expression. However, JDM is clinically heterogeneous, and is likely to have transcriptomic heterogeneity despite the presence of these general trends. We approached inter-individual variation of gene expression using Principal Components Analysis (**PCA**). We retained only genes with significant differential expression in the untreated vs. control contrast and a minimum 2-fold change. Plotting the top two PCs revealed several trends (**Fig. 8**). First, samples tended to cluster with their own group rather than being randomly interspersed. Second, the inactive samples are somewhat control-like, with some inactive samples clustering closely with the control group. But, importantly, they are for the most part still distinct. This is likely due to the persistent inflammatory signal in the inactive samples. Third, the untreated samples were quite heterogeneous. Some clustered near the control and inactive samples. Other untreated samples were far removed from them. This indicates that for the most significant differentially expressed genes that PBMCs from untreated individuals have substantial variability in the expression of the individual genes. This variability could be attributed to severity of disease, duration of untreated disease, choice of therapy, intrinsic genetic variation, or environmental factors.

**Figure 8.**
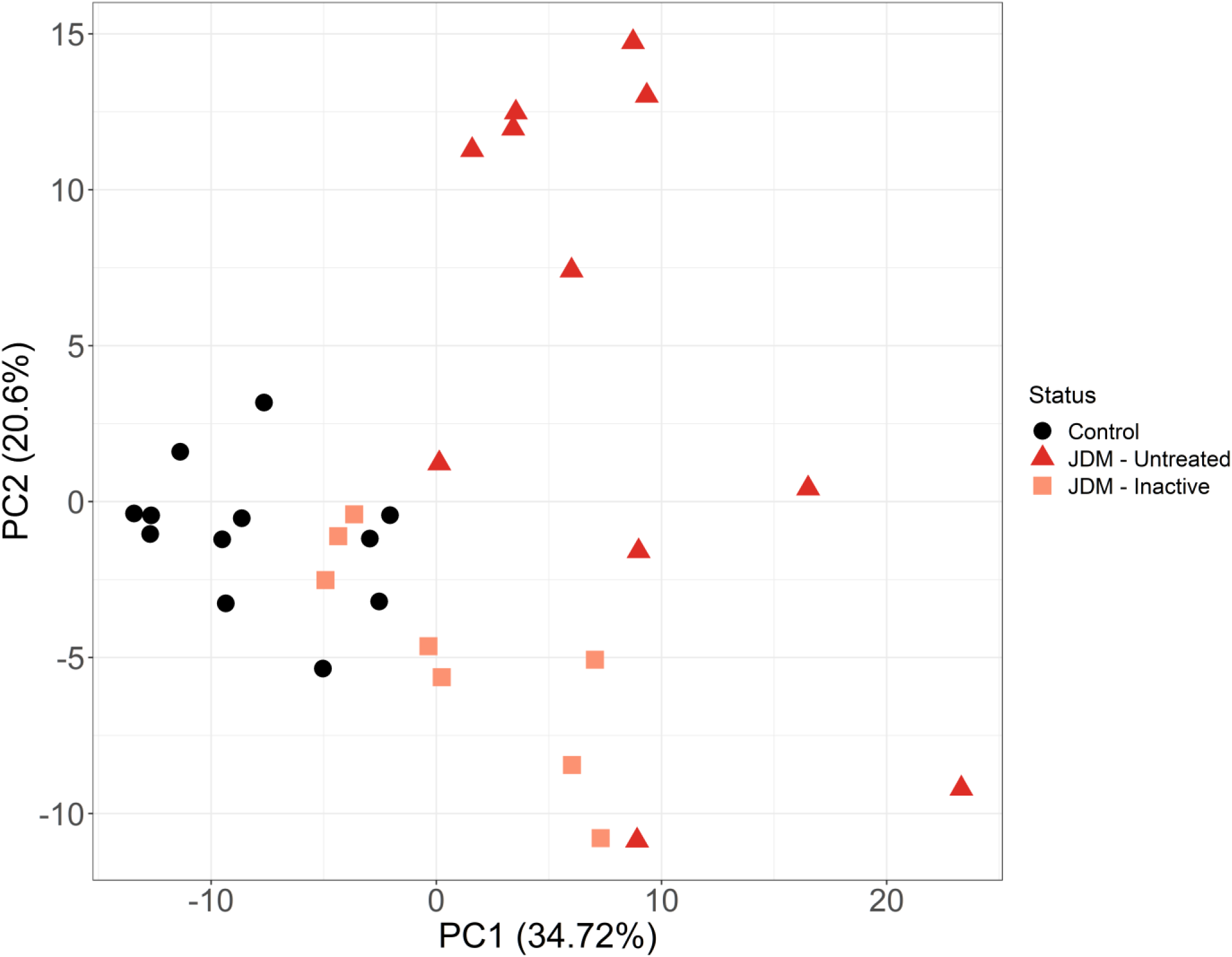
Principal components analysis shows substantial variability amongst untreated JDM samples and that inactive samples are mostly not control-like. Data from principal components analysis of JDM PBMC expression. The input data were regularized logarithm normalized transcriptome data. We only included genes with at least 2-fold differential expression (up or down) in untreated JDM vs. controls. The x- and y-axis are the first two principal components. Each dot is an individual sample, with disease status denoted by color and shape. Control PBMCs all cluster together. Inactive JDM samples range from control-like to distinct expression signatures. Some of the inactive samples are indistinguishable from untreated JDM. Perhaps most importantly, the untreated JDM samples show enormous variability even amongst each other, rather than clustering into a discrete, single group.

### 2.4. JDM skin has increased type 1 interferon and antigen presentation themes, with decreased cellular respiration and cholesterol synthesis

PBMCs are often a source of RNA for differential expression studies, as they are easy to access and low-risk. But while there is evidence of activation of circulating immune cells, inflamed tissue might be more informative in revealing the underlying molecular biology. We next tested for transcriptomic changes between samples from JDM skin (n=4) and skin from controls (n=5; **Table ST12**; **Fig. 9a**). There were 1,000 DEGs in this comparison, with 665 increased in JDM (656 at least 1.5-fold) and 335 decreased in JDM (323 at least −1.5-fold). The decreased genes included many ribosomal proteins (RP). Some of the most significant decreases were in *UQCRB* (−4.42 FC), *NGB* (−57.3 FC), and *FDPS* (−4.04 FC). *UQCRB* is part of mitochondrial complex III, and would therefore have obvious roles in cellular metabolism. It also has been shown to promote angiogenesis (23), which is interesting considering that the gene is decreased and that vascular damage is a central feature of JDM. As suggested by some of the top decreased genes, there were enrichments (**Table ST13; Fig. 9b**) for the ribosome (GO:0022626, GO:0022625, GO:0003735, GO:0044391), translation (REAC:R-HSA-156842, REAC:R-HSA-156902, REAC:R-HSA-72764), and protein targeting to the endoplasmic reticulum (GO:0070972, GO:0045047, GO:0072599) or membrane (GO:0006614, GO:0006613). A second major theme was cellular metabolism, specifically respiratory complex and oxidative phosphorylation (GO:0016627, GO:0006119, GO:0098803, GO:0070469). Decreased genes were also enriched for genes related to cholesterol synthesis (GO:0006695, GO:0016126, REAC:R-HSA-191273). Cholesterol plays a critical role in maintaining the integrity of the stratum corneum, and a decrease in genes related to this them may indicate that affected JDM skin is a more “leaky” barrier than control skin. Interestingly, despite the fact that these were skin samples, there was an enriched theme for muscle thin filament tropomyosin (GO:0005862) due to decreases in tropomyosin genes *TPM1*, *TPM2*, *TPM3*, and *TPM4*.

**Figure 9.**
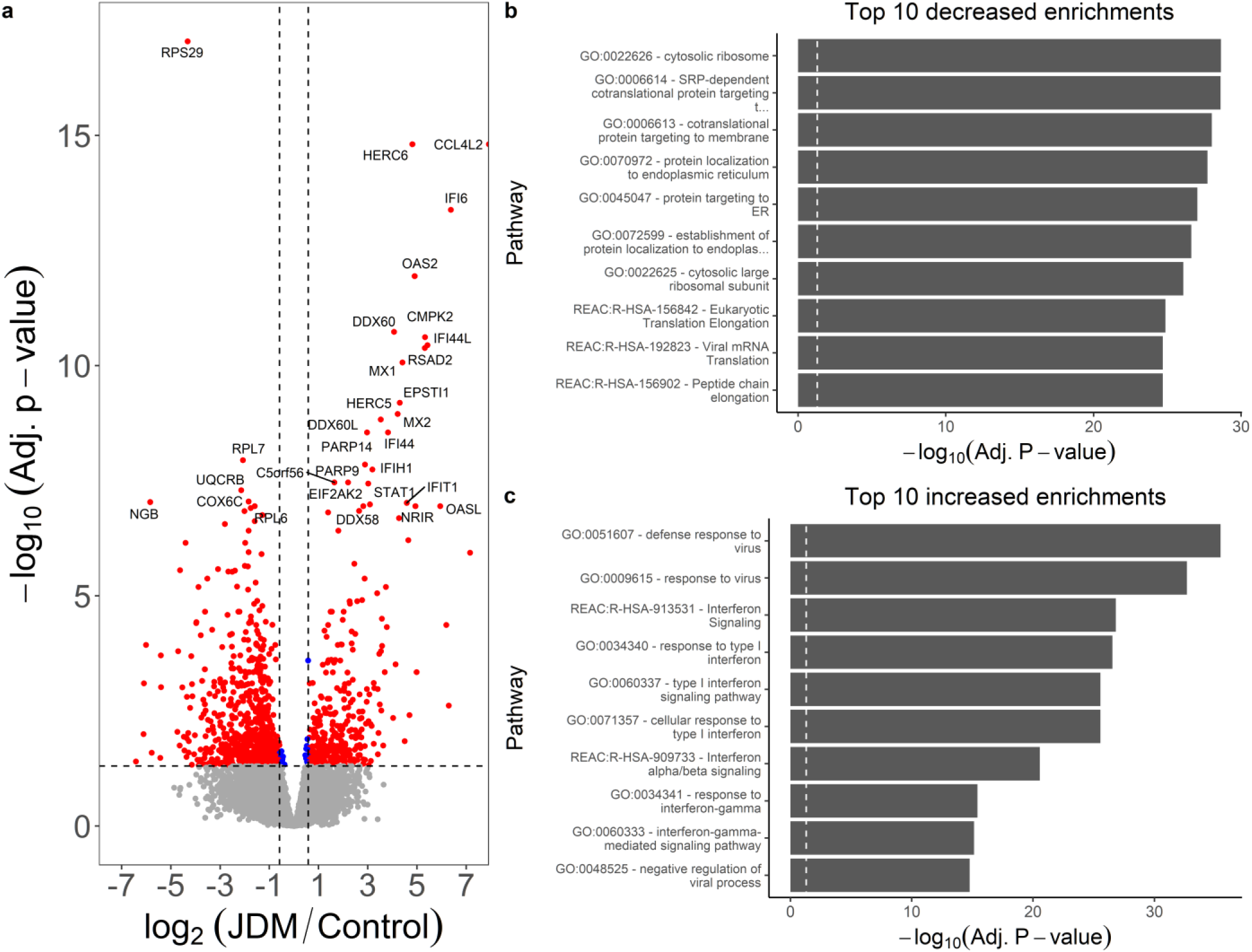
JDM skin has a strong signal for decreased translation and protein targeting, with enrichments for increased type 1 interferon signaling. **a.** JDM vs. control skin volcano plot. The x-axis is log_2_ fold-change and the y-axis is -log_10_ of false-discovery rate corrected p-value. The horizontal dashed line is the 0.05 significance cutoff, and the vertical dashed lines are minimum 1.5-fold changes. **b.** gProfileR enrichments for genes with decreased expression are enriched for translation and protein localization. **c.** Genes increased in JDM skin have a strong type 1 interferon signature.

Consistent with the DE genes in untreated JDM PBMCs, involved skin had increased inflammatory gene signatures and evidence for interferon-responsiveness. This included increased expression of *CCL4L2*, *IFI6*, *OAS2*, *IFI44L*, *MX1*, and *MX2*, among others. Several of the inflammation-related genes were nearly undetectable in the control biopsies, leading to improbably large fold-changes. Major themes for the genes with increased expression in JDM skin were overall also similar to untreated JDM PBMCs: enrichments for type 1 interferon signaling as well as for targets of STAT2 and the various IRF transcription factors (**Table ST14; Fig. 9c**). There were additional themes specifically for response to interferon gamma (GO:0034341, GO:0060333, GO:0071346, REAC:R-HSA-877300, GO:0072643), as well as antigen presentation (GO:0002483, GO:0019883, GO:0019885, GO:0002474). Consistent with MHC antigen presentation, *TAP1* was increased in skin (4.19 FC). TAP1 and TAP2 protein form the transporter associated with antigen processing protein complex. This raises the question of why *TAP1* is increased and why the themes are enriched. Particularly, is this the result of an increase in gene expression without a change in cell composition, or is it a result of the previously observed increased number antigen presenting cells in the skin of JDM patients (24)?

### 2.5. JDM muscle has decreased mitochondrial respiration, and increased interferon signaling and antigen presentation

We also tested for transcriptomic alterations in affected JDM muscle (n=4) compared to control muscle (n=5). This comparison resulted in the most extensive list of differentially expression genes, with a total of 1,697 significant genes (**Table ST15**; **Fig. 10a**). This included 957 genes with decreased expression (919 at least −1.5-fold) and 740 genes with increased expression (700 at least 1.5-fold). Among the decreased genes, there was some overlap with what was observed in skin. There were numerous top decreased genes that were associated with ribosomal function (such as the ribosomal RPS and RPL genes) or cellular respiration (such as *UQCRB*, *COX6C*, and *COX7C*).

Another top decreased gene was the long intergenic non-coding RNA *AL021920.3*. There is not any well-described function attributed to this RNA. While not encoding protein products, non-coding RNAs can have important regulatory functions both in cis and in trans, and may therefore affect the expression of many other genes. As might be expected, titin was significantly decreased (*TTN*; −2.63 FC). However, there was an even more significant decrease in the titin antisense transcript (*TTN-AS1*; −4.87 FC). This highlights the advantages of using a stranded RNA-Seq library preparation, as a non-stranded library would be unable to distinguish reads originating from an overlapping antisense transcript. Themes for decreased genes were overall similar to the themes for gene decreased in skin (**Fig. 10b**; **Table ST16**), such as splicing (GO:0071013, GO:0000375, GO:0008380, GO:0000398, REAC:R-HSA-156842), ribosome (GO:0022626), translation (REAC:R-HSA-156842, REAC:R-HSA-156902, REAC:R-HSA-72764), membrane targeting (GO:0006614, GO:0006613), and endoplasmic reticulum (GO:0045047, GO:0072599, GO:0070972). Mitochondrial respiration themes were again enriched (GO:0006119, GO:0098803, GO:0042775, GO:0098800, GO:0042773).

**Figure 10.**
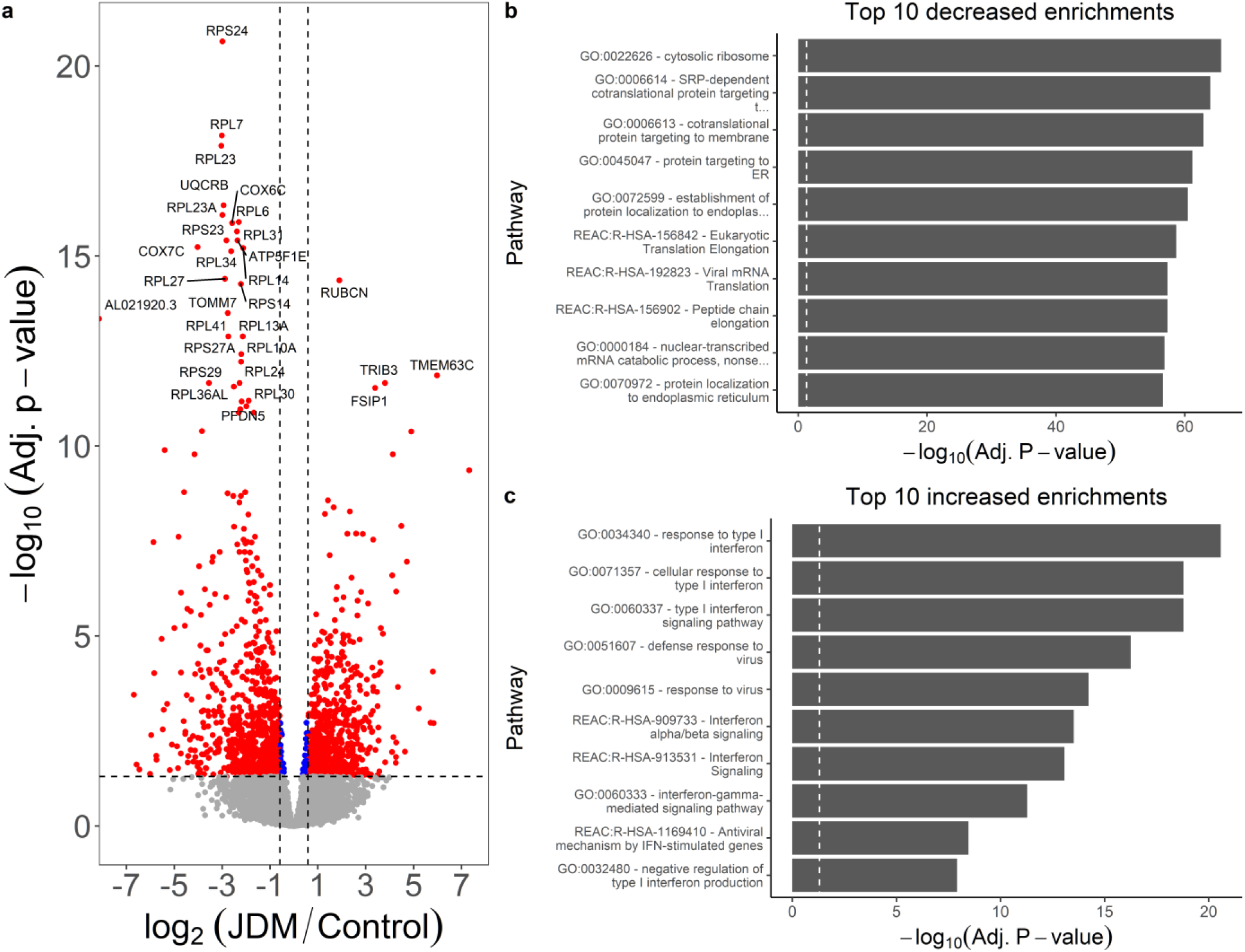
Genes decreased in JDM muscle are enriched for ribosome-associated proteins and translation, and increased genes are enriched for type 1 interferon response. **a.** Volcano plot of JDM muscle versus control muscle, with log2 fold-change (x-axis) and normalized expression (y-axis). Dashed lines indicate minimum 1.5-fold change (vertical) and adjusted 0.05 significance (horizontal) cutoffs. One of the top increased genes is *rubicon*, which acts to negatively regulate autophagy. This suggests autophagy could be an important player in JDM muscle. **b.** The top decreased genes are mainly ribosomal-associated proteins. Genes with decreased expression support this by showing enrichment for the ribosome, translation, and protein targeting. It’s unclear if these decreases reflect muscle degeneration. **c.** Genes with increased expression are enriched for type 1 interferon and interferon gamma signaling, concordant with PBMCs and skin samples from JDM.

With regard to genes with increased expression in JDM muscle, some of the most significant were *RUBCN* (3.17 FC), *TMEM63C* (62.91 FC), *TRIB3* (13.96 FC), *FSIP1* (10.50 FC), *IL31RA* (29.83 FC), and *LURAP1L-AS1* (17.49 FC). Rubicon (*RUBCN*) is part of the Beclin-1 complex, and acts to negatively regulate autophagy. It’s required for LC3-mediated phagocytosis, and also negatively regulates type 1 interferon signaling by preventing IRF3 transcription factor dimerization (25, 26). Interestingly, a quantitative trait locus study of >8,000 people found that SNPs in *TRIB3* are associated with the level of IL6 detectable in the blood (27). This suggests that *TRIB3* plays some role in regulating IL6 expression. IL31 signaling, including increased expression of IL31RA, is thought to have a role in the severity of itchiness in affected JDM skin (28). These results suggest there is increased IL31 signaling in affected muscle as well. As far as enriched processes, there was substantial overlap with skin and PBMC findings (**Table ST17**; **Fig. 10c**). A main signal was for both type 1 and type 2 interferon signaling (REAC:R-HSA-877300, GO:0034340, GO:0071357, GO:0060337, REAC:R-HSA-909733). Consistent with enrichments for interferon signaling, there was enrichment for targets of various interferon-responsive factor (**IRF**) transcription factors, as well as STAT2 (TF:M10080, TF:M09957, TF:M07216, TF:M11681, TF:M11671). There was again enrichment for antigen presentation through the MHC and TAP binding (GO:0046977, GO:0046979, GO:0002479, GO:0042590, GO:0042612, GO:0002483, GO:0042605).

With these comparisons completed, we could then examine the overlap between different tissues. We included differentially expressed genes in the untreated vs. control comparison, as well as the JDM vs. control comparisons for skin and muscle (**Fig. 11**). Overall, most differentially expressed genes were distinct for each tissue. The greatest number of differentially expressed genes was in muscle, followed by skin, followed by untreated PBMCs. The greatest overlap was between genes increased in both JDM skin and muscle (n=84) and decreased in both JDM skin and muscle (n=160). The next greatest overlap was genes increased in untreated PBMCs, skin, and muscle, primarily driven by increased interferon signaling, (**Table ST18**). This core set of increased genes included *ADAR*, *DDX60*, *DDX60L*, *IFI44*, *IFI44L*, *IFIT1-3*, *ISG15*, *OAS2-3,* and *STAT1*.

**Figure 11.**
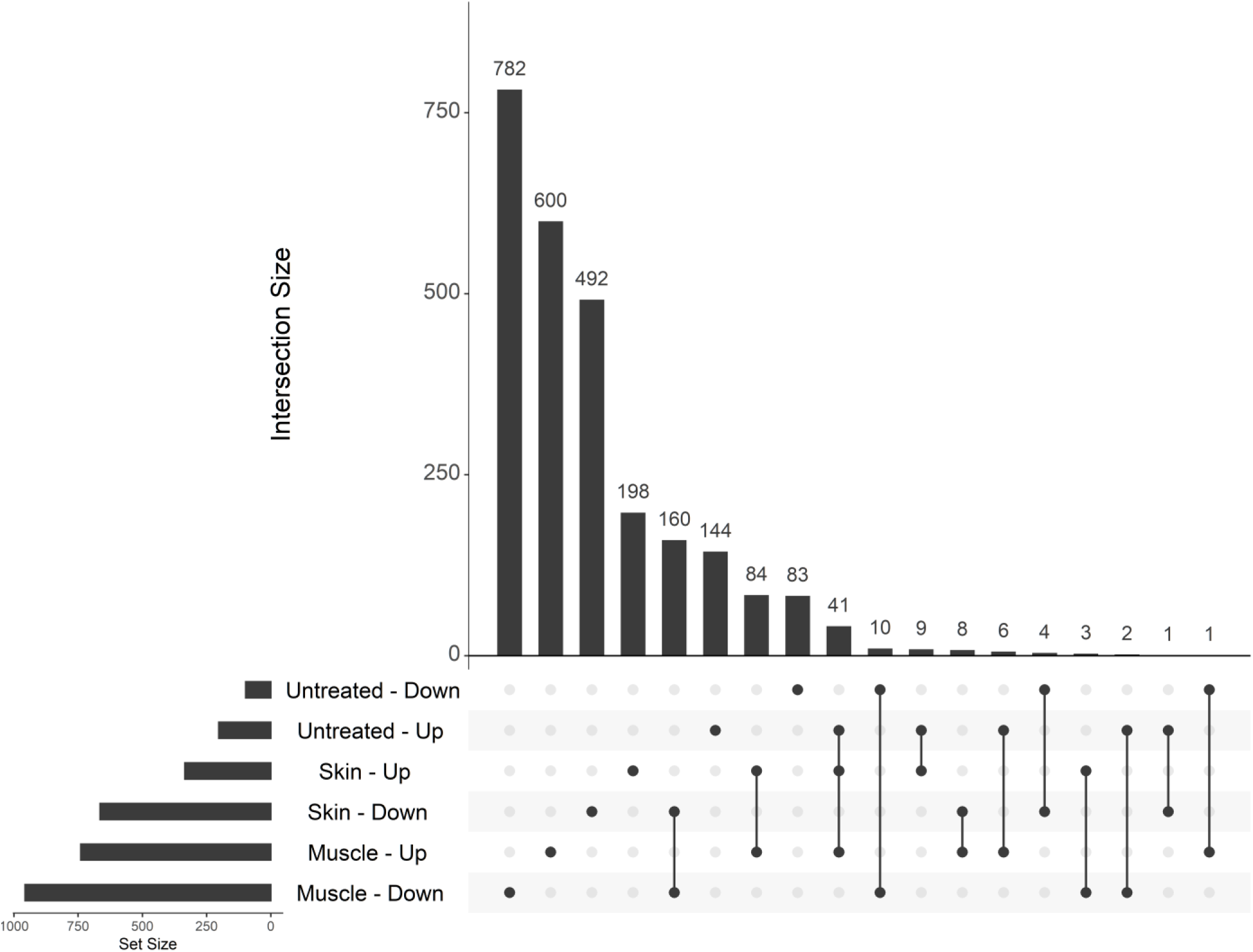
Most genes differentially expressed in JDM are specific to a given tissue, and overlapping genes between skin and muscle tend to have concordant directionality. Shown is an UpSet plot of the intersections between genes in the untreated JDM PBMC versus control comparison (Untreated), JDM skin versus control skin comparison (Skin), and JDM muscle versus control muscle comparison (Muscle). In each category the direction indicates whether a gene was increased (Up) or decreased (Down) in JDM. The bottom left bar plot shows the total number of DE genes in each category. The vertical bars indicate the number of genes in that particular intersection, which is defined by which dots are filled in. The greatest numbers of DE genes are in muscle, followed by skin. The greatest overlap is between skin and muscle decreased (n=160) and increased (n=84). Genes with concordant increases in JDM PBMCs, skin, and muscle are mainly type 1 interferon-responsive genes.

### 2.6. PBMC gene expression correlates with measures of JDM clinical severity

We tested for correlation normalized (variance stabilizing transform) expression between several clinical parameters (DAS-M, DAS-S, DAS-T, and nailfold capillary end row loop quantification [**ERL**]) using weighted gene co-expression network analysis [**WGCNA**] (29). We only included PBMC data from untreated and inactive samples. A summary of the number of negative and positive trait correlations is in **Table 4**. For DAS-M, there were 19 negative and 101 positive correlations in one expression module (**Table ST19**). Genes with negative correlations included *CASS4* (−0.70 correlation [**corr**]), *UBE2Q2P1* (−0.70 corr), *DSG2* (−0.67 corr), and *TOMM7* (−0.60 corr). There were no enriched themes for the negatively correlated genes. Top positive correlations included *TNFRSF19* (0.83 corr), *MANF* (0.80 corr), and *CHMP5* (0.79 corr). The genes that positively correlated with DAS-M were enriched for similar pathways related to type 1 interferon, RIG-I signaling (KEGG:04622, WP:WP3865), and protein polyubiquitination (GO:0000209; **Table ST20**).

**Table 4.** WGCNA network summary.

| **Trait** | **Negative** | **Positive** |
| --- | --- | --- |
| DAS-M | 19 | 101 |
| DAS-S | 56 | 111 |
| DAS-T | 81 | 351 |
| ERL | 393 | 26 |

Summary data for the weight-gene co-expression network analysis (**WGCNA**). Four clinical traits were tested: Disease Activity Score for muscle (**DAS-M**), skin (**DAS-S**), and total activity (**DAS-T**), as well as quantification of nailfold capillary end row loop quantification (**ERL**). For all DAS, there were more positive correlations than negative correlations. For ERL, there were more negatively correlated genes. The fewest correlations were with DAS-M, and the greatest number of correlations was with DAS-T.

DAS-S had 56 negative and 111 positive significant correlations with gene expression in 3 separate expression modules (**Table ST21**). There were many correlations with ribosomal proteins (RPS and RPL genes). Other negatively correlated genes included *PNN* (−0.74 corr), *MALAT1* (−0.71 corr), *NSA2* (−0.71), and *GMFG* (−0.68). *MALAT1* is an important regulator of interferon responsiveness. In the mouse RAW264.7 macrophage-like cell line, *Malat1* is decreased after vesicular stomatitis virus infection, and deletion of *Malat1* leads to increased production of both IFN-α and IFN-β (30). The negative correlation with *MALAT1* suggests that it is a proxy for activation along the type 1 interferon axis. The major pathway enrichments were related to ribosomal function and protein targeting (**Table ST22**). Positive correlations with DAS-S included *CCL2* (0.78 corr), *NDN* (0.77 corr), *AC025524.2* (0.77 corr), and *KIF15* (0.71 corr). In mice, *Ccl2* expression depends on *Ifnar*, and it helps recruit both T cells and natural killer cells to sites of viral infection (31). The strongest correlation might therefore represent a strong signal for attracting immune cells to the skin. The genes positively correlated with DAS-S were primarily type 1 interferon-responsive genes (**Table ST23**).

Total DAS score is calculated by summing the muscle and skin scores. One might therefore expect the correlation and associated pathways to reflect what was seen in the individual DAS-S and DAS-M correlations. The DAS-T significantly correlated genes (n=432 in 5 modules) do indeed reflect this trend (**Table ST24**). Genes with the most significant negative correlations with DAS-T include *RPLP1* (−0.82 corr), *UBE2Q2P1* (−0.75 corr), *TOMM7* (−0.68 corr), and *MALAT1* (−0.62 corr). The major pathway signals included ribosomal categories, translation, and nonsense-mediated decay (**Table ST25**). The genes positively correlated with DAS-T were primarily interferon-related genes, such as *IFI6* (0.91 corr), *RSAD2* (0.90 corr), *IFI44* (0.90 corr), and *MX1* (0.88). The pathways were also consistent and primarily reflected the interferon signature (**Table ST26**).

The nailfold capillary end row loop correlations were mostly different from the correlations with clinical disease activity scores. For DAS, an increasing score reflects greater disease activity. For ERL, increased vascular damage would lead to a lower ERL, so some genes correlated with different DAS domains might be inversely correlated with ERL. Another clear difference is the category of correlated genes. Overall 63% of genes correlated with ERL are pseudogenes (**Table ST27**). It is unclear why this particular category is overrepresented for ERL. In total, there were 419 genes that correlated with ERL in 2 different modules. Among protein coding, antisense, and non-coding RNAs, the most significant negatively correlated genes included *RAB6C* (−0.81 corr), *EIF5AL1* (−0.81 corr), and *PABPC3* (−0.77 corr). The large number of pseudogene correlations made it difficult to assign the genes to known pathways. There were only 4 enrichments, all of which were related to type 1 interferon (**Table ST28**). There were fewer genes (n=26) that were positively correlated with ERL. There were some pseudogenes in this set, but they didn’t predominate. Top positively correlated genes included *RNF216* (0.63 corr), *TBX18* (0.63 corr), *GPATCH8* (0.62 corr), *NRF1* (0.57 corr), and *GTF2E2* (0.57 corr). The only pathway enrichment was for targets of microRNA hsa-miR-33a-3p (**Table ST29**).

After having identified genes correlated with different clinical parameters, we could then determine if these genes are just a subset of differentially expressed genes (**Fig. 12**). The greatest set were genes were only differentially expressed and not correlated with any clinical trait (n=236). The next greatest set were genes that correlated with both end row loop number and total DAS score (n=215), followed by genes correlated to ERL only (n=159). Looking across differentially expressed genes and trait-correlated genes, 78% of differentially expressed genes in untreated JDM did not correlate with any trait, and 89% of trait-correlated genes were not differentially expressed. This is a key point, as it suggests that while case-control differential expression studies can help us understand the underlying disease molecular biology, it would be a poor method to identify biomarkers of disease activity. There were 21 genes that were both correlated with every trait and differentially expressed, including *AGRN*, *CCL2*, *CXCL10*, *IFI27*, *IFIH1*, *IFIT1-3*, *MX1*, and *TRIM22* (**Table ST30**).

**Figure 12.**
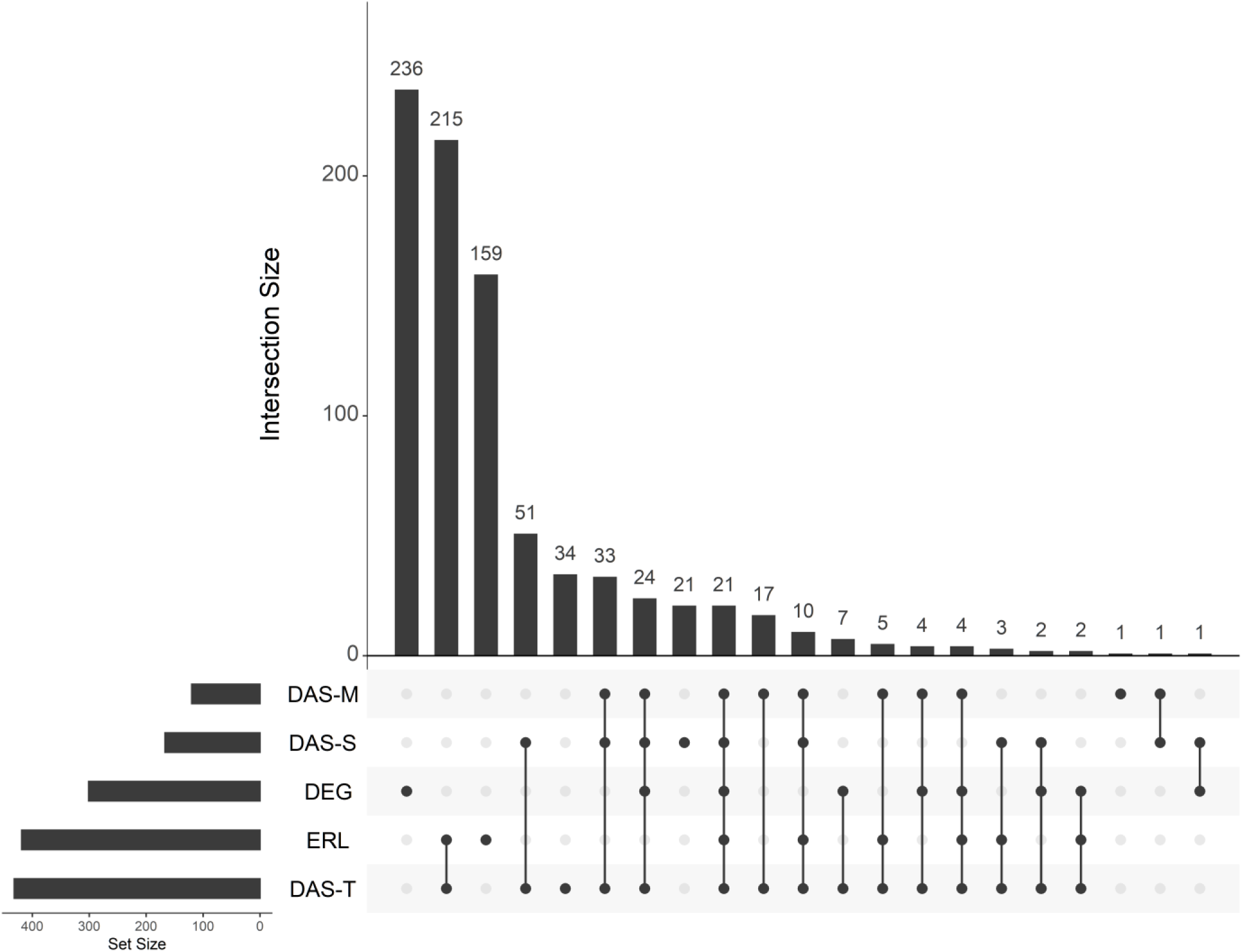
Genes associated with disease severity measures are not typically differentially expressed. An UpSet plot of intersections between untreated JDM vs. control PBMCs (DEG) and genes correlated with 4 disease severity clinical traits. The lower left bar plots show the total number of genes in each category. The vertical bar plot shows the intersection size. Intersections are defined by which dots are filled in and connected. Most of the differentially expressed genes are not associated with traits, and most trait-associated genes are not differentially expressed. The greatest intersection is between genes correlated with total DAS and nailfold capillary end row loop number. As far as differentially expressed genes go, the greatest overlap is a set of 24 genes that are differentially expressed and correlated with all three DAS, followed by a set of 21 genes that are differentially expressed and correlated with every clinical trait. Abbreviations: DEG, differentially expressed genes; DAS, disease activity score; ERL, nailfold capillary end row loop number.

We then wanted to examine the gene-gene correlation network amongst all genes that correlated with any trait. We took the biweight midcorrelation for all pairs of genes in this, retaining any gene pair with at least 80% correlation. We then imported the genes as nodes and the gene-gene correlation as edges in Gephi. This allowed us to both visualize the gene-gene correlation network and calculate relevant network statistics (**Fig. 13**). We ranked genes by weighted degree and page rank to identify genes that had central roles in the network (**Table ST31**). The 22 highest ranked genes all correlated with DAS-M, DAS-S, and DAS-T. Among the top ranked genes were *RSAD2* (weighted degree = 80), *IFIT1* (weighted degree = 80), *CMPK2* (weighted degree = 78), *PARP9* (weighted degree = 76), and *TRIM22* (weighted degree = 75). Both *LY6E* and the non-coding *LY6E-DT* divergent transcript were highly ranked, with weighted degrees of 66 and 61, respectively. Otoferlin (*OTOF*) was also correlated with all 3 DAS domains (weighted degree = 43). Some of these genes that correlated well with all DAS scores and are highly connected, central genes in the network could be candidates for a more quantitative, molecular measure of disease activity score. It is, however, still noteworthy that these markers correlate with DAS, and therefore don’t capture the genes that continue to be differentially expressed when the child appears to be clinically inactive.

**Figure 13.**
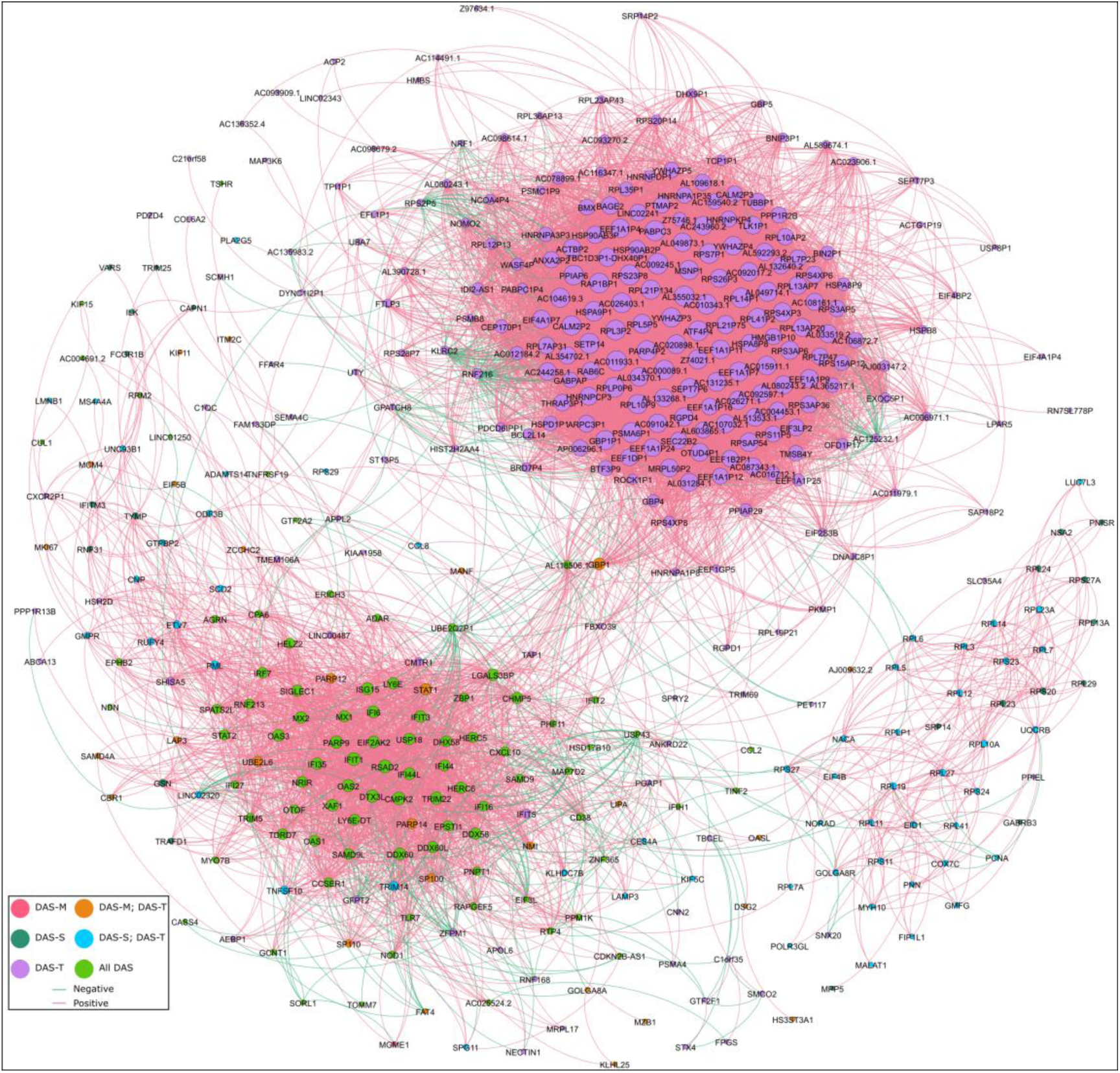
Gene-gene correlation analysis of trait-correlated genes shows a sub-network of genes correlated with DAS-T only and another correlated with all DAS. Network representation of all DAS-correlated genes that have at least 0.80 pairwise gene-gene correlations. Each node is an individual gene in the network. The node color is determined by which traits the gene is correlated with. The node size is partitioned by weighted degree. Each edge is colored based on whether the gene-gene correlation is positive or negative. Most of the highly interconnected genes are positively correlated with each other, though this is not always the case. There are two main sub-networks that are apparent. The lower left network is composed mainly of genes correlated with all DAS, and is mainly composed of type 1 interferon-responsive genes (*RSAD2*, *IFIT1*, *MX1*, *PARP9*, etc). The other main sub-network is at the upper right quadrant, and is composed mainly of genes only correlated with total DAS. This network is composed more of genes related to translation (*EEF1A1P11*, *EEF1A1P7*, *EEF1A1P9*, etc) and ribosomal proteins (*RPL41P2*, *RPL3P2*, *RPL5P5*, etc).

### 2.7. Comparison of expression to published SOMA protein data

We had previously used the SOMA protein array technology to test for differences in plasma protein levels of untreated JDM versus controls and whether the protein levels change in response to treatment (32). We wanted to compare the SOMA plasma results to the PBMC RNA results of this study. The same patients and time points were not used between the two studies, so we compared summary statistics. The relationship between the technologies wasn’t 1:1. The SOMA targets sometimes correspond to more than one Uniprot ID, depending on whether the platform probes a protein or complex. Conversely, some individual genes were represented by multiple Uniprot IDs on the SOMA platform. We merged the datasets by listing one SOMA Uniprot ID and one Ensembl gene ID per line. First, we examined the results of untreated JDM versus healthy controls. Overall, 209 of 251 differentially expressed SOMA targets representing 218 Uniprot IDs mapped to 219 Ensembl gene IDs (**Table ST32**). Only 12 targets had differentially expressed RNA in PBMCs and differentially detected protein on the SOMA. There were 3 with discordant fold-changes (*ADAM12*, *CFH*, *FGF2*), and 9 with concordant fold-changes (*CCL2*, *CCL3L1*, *CXCL1*, *CXCL10*, *EPHB2*, *IL1RN*, *ISG15*, *LGALS3BP*, *STAT1*) between the plasma protein and PBMC RNA. We then compared the results of treatment for 11/12 of these targets (**Table ST33**). There was agreement in treatment response for 7 targets: 3 where the treatment led to improvement of the initial dysregulation (*CXCL10*, *ISG15*, *LGALS3BP*) and 4 where treatment failed to resolve the dysregulation (*CCL3L1*, *CFH*, *CXCL1*, *IL1RN*). The other 4 targets had disagreeing responses to treatment with the two technologies (*ADAM12*, *EPHB2*, *FGF2*, *STAT1*). Given the limited overlap between the two approaches, these technologies are complementary to each other rather than being redundant.

## 3. Discussion

Prior to the use of corticosteroids, the mortality rate for JDM was as high was 1 in 3 (33). With the introduction of corticosteroids, the mortality rate has improved to less than 2% in the US and United Kingdom. Despite these treatment advances and improved outcomes, there is still significant morbidity associated with JDM, such as pulmonary compromise (34) and the development of calcinosis, sometimes with superinfection (5). Inpatients with a diagnosis of JDM have increased rates of hypertension, atherosclerosis and lipodystrophy (35). Additional evidence suggests that adults who had JDM as children have an increased risk of hypertension and lipodystrophy, along with increase endothelial intima media thickness (8). These data together suggest that while treatments have improved outcomes, there is more work to be done to reduce the morbidity associated with JDM. This is an important issue, as few adults who had childhood JDM seek assessment for increased cardiovascular disease risk. We sought to better understand JDM by performing RNA-Seq on untreated and control JDM PBMCs. We also characterized clinically inactive samples from a subset of the same individuals with JDM to determine if they had reverted to a control-like state. Separately, we examined skin and muscle from JDM and control individuals to determine how well PBMCs reflect the types of changes observed in directly affected tissues.

One surprising finding was reduced expression in JDM patients of several genes associated with autophagy (*ST8SIA* and *ARL3* in untreated PBMCs; *RUBCN* in muscle). Autophagy is a critical cellular process that involves the catabolism of cell components and organelles, which has previously been linked to some autoimmune diseases (36). Reduced autophagy has been previously implicated in systemic lupus erythematosus, psoriasis, rheumatoid arthritis, and inflammatory bowel disease (37). Reduced autophagy can lead to an increased presence of old, damaged cellular components, which can elicit immune responses and antibody generation. It is possible that reduced autophagy plays a role in the initiation of JDM as well.

Another unexpected finding was increase expression of *otoferlin* in untreated JDM. Mutations in another ferlin family member, *dysferlin*, were initially found to be associated with Limb Girdle Muscular Dystrophy type 2B and Miyoshi myopathy (38). Mutations in *otoferlin*, on the other hand, are associated with non-syndromic deafness (39). It also likely plays a role as a calcium-sensitive scaffolding protein that can interact with SNAREs (40). It’s therefore possible that in JDM that *otoferlin* may have a role in immune cell calcium flux, and perhaps calcium-dependent granule discharge.

Previous studies identified increased type 1 interferon signaling in JDM PBMCs (41, 42). We confirmed this finding along with some additional observations. First, while there is a clear type 1 interferon signature, there is extensive inter-individual variation in the expression levels of interferon-responsive genes in untreated JDM patients. This is likely mediated by disease severity, duration of untreated disease, environmental factors, and genetic differences, but the dynamic interaction of these different variables remains to be evaluated. Second, while type 1 interferon predominates, there is also evidence for IL-1, IL-10, and NF-κB signaling. This highlights the fact that the immune response in JDM is complex and involves many different signaling pathways. A third observation is related to treatment response. JDM patients are typically treated until they reach accepted criteria for clinical inactivity (43, 44). We observed that patients who appeared to be clinically inactive had a robust immune activation signature. Part of this signature was related to type 1 interferon response, but the major signal was for IL-1, with some evidence for IL-10 and NF-κB signaling.

We compared the transcriptional results from untreated JDM PBMCs to the data we published from a previous paper examining sera proteins in untreated JDM as well (32). Overall only 12 protein targets were altered in untreated sera and also differentially expressed in untreated PBMCs. This finding highlights that serum protein and PBMC transcriptomes are not a substitute for one another, and that each may have unique predictive power in the clinic.

Examining affected JDM skin and muscle confirmed that the increased interferon signature is present in both tissues, along with a concurrent enrichment for translation and the ribosome amongst genes with decreased expression. This is consistent with previous results that showed increased type 1 interferon in JDM muscle when compared to healthy controls (45). Another study showed that JDM muscle biopsies had increased expression of OAS family genes, specifically *OAS1*, *OAS2*, *OAS3*, and *OAS4* (46). The authors were particularly interested in the OAS family of genes because they are important during infections with double-stranded RNA viruses. The potential association of JDM onset with preceding infection with an RNA virus such as Coxsackievirus B2 or B4 has long been a topic of interest (47, 48). We can confirm increased expression of *OASL*, *OAS1*, *OAS2*, and *OAS3* in JDM skin, muscle, and PBMCs.

We also found evidence for mitochondrial dysfunction in both skin and muscle, as both had enrichments for terms involving oxidative phosphorylation, mitochondria membrane, and the respiratory chain. Mitochondrial dysfunction and reduced expression of mitochondrial genes has been previously observed in muscle biopsies from adult dermatomyositis patients (49).The same study documented that treating human myotubes with IFN-B significantly reduced their mitochondrial respiration, suggesting a link between type 1 interferon and mitochondrial dysfunction. Our findings suggest that this dysfunction could also occur in affected skin.

There is some additional data available from adult dermatomyositis. RNA-Seq transcriptome comparison of polymyositis and dermatomyositis has shown dermatomyositis-specific increases in *OASL*, *EPSTI1*, and *IFI27* within CD8+ T cells (50). *OASL* and *EPSTI1* were increased in skin, muscle, and untreated PBMCs in our study. Another study used Agilent microarrays to analyze the transcriptome of muscle biopsies from 3 adult dermatomyositis patients and 5 controls (51). The muscle showed an increase in the gene *MX1*, but its gene ontologies were not as enriched for interferon specifically. Most of the top pathways were related to general increased inflammation and leukocyte activation. A third study in adults looked at gene expression with Affymetrix arrays using muscle tissue from controls (n=5) and dermatomyositis (n=5) (52). Their findings agree with the interferon signature, but interestingly also pointed to increases in *DDX58*, *DDX60*, and *IFIH1* that suggested the RIG-I signaling pathway (DDX58 is the RIG-I gene). *DDX58* was not significantly different in untreated vs. control PBMCs, but did appear to be elevated in the untreated vs. inactive comparison, likely due to better power for the paired test. In contrast, in this study *DDX58* was increased in JDM skin (7.06 FC) and muscle (5.06 FC). Since our data show that the increased RIG-I expression resolves with treatment, this particular pathway may be more associated with overtly active disease.

In both skin and muscle we also saw enrichments for MHC class I antigen presentation, associated with increases in *HLA-B*, *HLA-C*, *HLA-F*, *HLA-G*, *TAP1*, and *TAP1BP*. Increased MHC I protein has been observed in JDM muscle biopsies (53, 54). Similar to many other autoimmune and inflammatory diseases, the primary signal in genome-wide association studies for adult and juvenile dermatomyositis is in the major histocompatibility complex (55). If this up-regulated MHC class I expression is stimulating a sustained immune response, the specific antigens have not yet been identified, but are under active investigation.

We performed a trait-gene correlation analysis along with a gene-gene correlation analysis of trait-correlated genes to identify potential molecular biomarkers of disease activity. One motivation for this was that while JDM clinical disease severity scores are generally considered reliable, there is both inter-rater variability, which obviously complicates multi-center research and clinical trials. Molecular biomarkers of disease activity have the potential to be more sensitive, less variable, and more scalable than physical assessments. The trait-correlated genes themselves may prove useful as molecular biomarkers for different disease domains, and the network analysis provides guidance for “hub” genes that are both correlated with traits and highly connected in the gene-gene correlation network. One of these hubs is *RSAD2*, which is also increased in primary Sjogren’s syndrome (56) and psoriasis (57). *RSAD2* was also the most highly ranked hub gene in a study to identify shared molecular etiologies for pemphigus and systemic lupus erythematosus using WGCNA analysis (58). Another central gene is *PARP9*, alternatively referred to as *BAL1* or *ARTD9*. It promotes the response to interferon in primary macrophages (59). Neopterin, a macrophage-derived product, is elevated in both untreated JDM and during flares (60). This suggests an important role for macrophages in the interferon-driven response of JDM.

This study relied on retrospectively collected PBMCs and tissue, and therefore has limitations that are important to recognize. There were limited numbers of PBMC samples available from treatment naïve patients, which correspondingly decreases our power and ability to assess how heterogeneous the transcriptomes are in untreated samples. The skin and muscle samples, while snap frozen, were not RNA stabilized. An optimum future design might involve a multi-center, prospective, longitudinal collection of samples stored specifically for high-throughput methodologies. This would enable a large enough cohort to specifically account for the effects of different treatment regimens. Perhaps most importantly, study of longitudinal samples will be required to ascertain what fraction of clinically inactive patients have chronic, smoldering tissue inflammation, and to determine how long this inflammatory state persists.

## 4. Methods

### 4.1. JDM patient population ascertainment and diagnostics

All individuals with JDM had an MRI (T2 weighted image) compatible with the diagnosis of an inflammatory myopathy. MSAs were determined on patient sera by immunodiffusion or immunoprecipitation at the Oklahoma Medical Research Foundation Clinical Immunology Laboratory (61).

A pediatric rheumatologist assessed the child at each visit to determine their disease activity score (**DAS**). A score of zero indicates no activity, and higher scores indicate greater activity. The skin domain (**DAS-S**) score ranges from 0-9, muscle domain (**DAS-M**) ranges from 0-11, and the total DAS (**DAS-T**) ranges from 0-20 (10). The team’s experienced physical therapists obtained childhood myositis assessment scores (**CMAS**), which range from 0 (worst) to 52 (best), with 52 indicating full strength and endurance for patients (62). Similarly, for healthy children age 4, a standard of 46 was established (63).

For end row loop (**ERL**) quantification, images of each of the eight digits, excluding thumbs, were captured and recorded for analysis later. The majority of ERL images were taken within 48 hours of sample collection. One untreated JDM sample was taken 24 days prior to sample collection when the individual was still untreated. Earlier photos were taken using freeze-frame video microscopy (12x) and printed in real-time on photo paper, where it was analyzed. Later, digital images were obtained via a Dermlite II ProHR (18x) with Nikon camera adapter, and analysis was performed utilizing Photoshop CS5; the two methods are concordant (64). After standardizing for magnification, ERL/mm was quantified by counting the number of end row capillary loops per 3mm section on each of the eight fingers, dividing this by three, transforming this count into ERL/mm. The representative ERL/mm for each patient was the average of all eight digits as we’ve previously reported (7, 65).

### 4.2. RNA isolation, sample storage, and library preparations

PBMCs were pelleted by Ficoll gradient density centrifugation of whole blood within 2 hours of acquisition. The pellets were resuspended in Recovery™ Cell Culture Freezing Medium (ThermoFisher #12648010) and stored in liquid nitrogen until RNA extraction. After allowing the frozen aliquots to thaw, the cells were collected with low-speed centrifugation (5’ at 500 xɡ), the supernatant aspirated, and 700 µL of Qiazol added directly to the pellet. For muscle and skin, the frozen samples were first stabilized with RNAlater ICE (ThermoFIsher AM7030). They were then finely minced and 700 µL of Qiazol was added.

All samples were thoroughly vortexed to facilitate cell lysis and the slurry was passed through Qiashredder columns (Qiagen #79654) for homogenization. Total RNA was isolated using the miRNeasy mini kit (Qiagen #217004) according to the manufacturer’s instructions, and the isolated RNA quantified using a Qubit 3 with the Qubit RNA HS kit (Invitrogen #Q32852). Stranded, total RNA libraries were generated using a ribosomal depletion method with template switching (Takara SMARTer Stranded High-Input Mammalian Total RNA Sample Prep Kit; #634875) according to the manufacturer’s instructions with limited modification. The first-strand synthesis incubation was extended to 2 hours to increase yield. During bead size selection, the beads were resuspended by pipetting with a narrow 1-20 µL tip. The beads were ethanol washed once instead of twice. For DNA elution from beads, incubation time was increased to 30 minutes, and the eluted DNA was moved to a new tube for PCR (14 cycles). Each sample used a unique combination of i5 and i7 indexes (combinatorial indexing). We combined the individually indexed libraries into variably-sized pools for sequencing. Each library pool was sequenced on one or more sequencing runs as needed to increase overall sequencing depth. We used low-coverage RNA-Seq (**Table ST1**; average 6.9M read-pairs / sample) to estimate the abundance of each gene. The pools were sequenced in paired-end mode with either 101 (HiSeq2500) or 150 (HiSeq3000) basepair cycles.

### 4.3. Processing sequencing data and calculating differential expression

The raw data (FASTQ) was cleaned by trimming sequencing adapters and low-quality terminal bases with cutadapt v1.15 (−-cut 6, -q 10, -m 30, -O 5, -a AGATCGGAAGAGC, -A SSSAGATCGGAAGAGC) (66). The cut and “SSS” flag on the adapter are because the kit uses Moloney Murine Leukemia Virus reverse transcriptase that adds a non-template triple C at the end of the first strand. These flags remove the non-template sequence. The cleaned data were aligned to Ensembl’s GRCh38 human reference genome using RNA-STAR v2.6.0c (67), and the number of reads for each annotated gene counted using featureCounts in stranded mode v1.6.3 (68). We counted all reads that overlapped with a known gene on the correct strand since we expected both nascent and spliced transcripts.

DESeq2 (v1.24.0) was used to calculate differential expression after summing the total read-pairs per gene for each biological sample (69). It was also used to calculate normalized expression with the regularized logarithm and variance stabilizing transform. For unpaired, two-group tests, sex was accounted for by including it as a covariate in the DESeq2 model (Wald test). This appropriately mitigated sex bias, as the reciprocal test (test for sex differences adjusting for disease status) identifies genes with known sex-biased expression, such as *XIST* (extended discussion in supplement). For paired samples, an individual identifier was included in the DESeq2 model to account for the pairing. We considered a gene to be differentially expressed if the false-discovery rate corrected p-value was less than 0.05 with no minimum fold-change.

We used the gProfileR tool (g:GOSt) to test for enrichments in known pathways, including only significant genes with a minimum 1.5-fold change. We always tested genes with increased expression separately from those with decreased expression using the following options: significant only, ordered query, no electronic GO annotations, hierarchical sorting, moderate parent term filtering, and 500 max size of functional category. Figure generation and statistical analysis was performed in R (v3.6.0).

### 4.4. Gene-network analysis

We calculated the correlation of the ‘vst’ normalized JDM PBMC (untreated and inactive) sample gene expression with four clinical parameters (DAS-S, DAS-M, DAS-T, and ERL) using the weighted-gene co-expression network analysis (**WGCNA**) package (v1.68). For any steps that involved calculating correlation we used biweight midcorrelation rather than Pearson correlation. Since there are several steps that involve matrix math operations in this type of network analysis, we used the Intel Math Kernel Library to speed the relevant operations. This identified genes correlated with the traits of interest. We then took all correlated genes and calculated all gene-gene pairwise midweight bicorrelations, with a minimum absolute correlation cutoff of 0.80. We used Gephi (v0.9.2) to visualize the network and calculate relevant network statistics (Modularity, Degree, PageRank, etc).

### 4.5. Validation with RT-qPCR

Reverse-transcription quantitative PCR (**RT-qPCR**) was used for otoferlin validation. A total of 600 ng of RNA was reverse-transcribed with random hexamers using the SuperScript first-strand synthesis kit according to the manufacturer’s instructions (Invitrogen, U.S.A). The cDNA (30 ng) was subsequently used in a 20 µL reaction with TaqMan Universal PCR Master Mix with an otoferlin TaqMan assay (Hs00191271_m1) or beta-actin reference gene (Hs01060665_g1). The experiments were performed in a 7500 Fast Real-Time instrument (Applied Biosystems) using Standard Mode cycling conditions. The significance of all pair-wise comparisons with multiple test correction was determined by the Tukey honest simple differences test in R.

### 4.6. Availability of data and material

These samples were from a retrospective collection that did not include approval for broad dissemination of sequencing data. However, the gene counts, which were used for the entirety of the analysis and are not identifiable, have been deposited in FigShare.

FigShare project: https://figshare.com/projects/2019_Untreated_juvenile_dermatomyositis_JDM_RNA-Seq/63539

Associated code (GitHub): https://github.com/RobersonLab/2019_UntreatedJDM_RNASeq

### 4.7. Study approval

All listed IRB approved protocols used the same age requirements. For ages 0 – 17, we obtained written informed consent from the participant’s legal representative. For ages 12 – 17, we also obtained the assent of the participant. Age-appropriate written informed consents / assents were used in each case.

All study protocols were approved by the Ann & Robert H. Lurie Children’s Hospital of Chicago IRB committee (IRB #2008-13457). For control samples, the study coordinator screened the volunteers to confirm they had no confounding medical illnesses or current prescription drug usage. They were then enrolled in the study using age-appropriate control consent (IRB #2001-11715). Only individuals with a clinical diagnosis of definite or probable JDM were eligible for inclusion in the JDM category (4). Interested individuals that met these criteria were enrolled for sample collection with biorepository tracking (IRB #2010-14117) and inclusion in our JDM registry of >580 children with confirmed JDM. All research was conducted in accordance with these approvals and was consistent with the Declaration of Helsinki.

## Author contributions

LMP and EDOR developed the idea for the study. GAM managed the clinical and sample databases, generating the relevant demographic and treatment tables. RAM and LC isolated RNA and generated sequencing libraries. WM performed RT-qPCR experiments. EDOR analyzed the RNA-Seq data and drafted the manuscript. All authors read, revised, and approved the manuscript.

## Acknowledgments

The authors would like to thank the patients who participated in this study, as well as the other physicians who helped care for these patients (K. Ardalan and M.L. Curran). We also thank Maria Amoruso for help with control recruitment. We also thank the Department of Pediatric Orthopedics at the Ann & Robert H. Lurie Children’s Hospital of Chicago for their assistance in obtaining control skin and muscle. This work was supported in part by NIH grants R21-AR066846 (LMP), P30-AR048335 (EDOR, LC) and P30-AR073752 (EDOR), as well as a grant from the Cure JM Foundation (LMP). The Genome Technology Access Center at Washington University generated the sequencing data, and is supported by Siteman Cancer Center grant P30CA91842 and ICTS/CTSA grant UL1TR000448. Some analyses were performed on the Center for High Performance Computing (CHPC) cluster at Washington University, which is partially supported by NIH grant S10 OD018091.

