## Supplemental methods & information for "Transcriptomes of peripheral blood mononuclear cells from juvenile dermatomyositis patients show elevated inflammation even when clinically inactive"

Elisha D.O. Roberson, Ph.D.
Washington University in St. Louis
660 South Euclid Avenue
Campus Box 8045
St. Louis, MO 63110
eroberson [AT] wustl [DOT] edu

---

**Supplementary Material**

**6. Supplementary table legends**

Read-pairs per sample – ST1
RGSM: Read Group Sample ID.
Status: Disease status.
Sex: F – female; M – male.
Tissue: Source of tissue.
Total: Total read-pairs for sample.
Additional columns: Individual sample run read-pair count.

Inactive time point medications – ST2
RGSM: Read Group Sample ID.
Meds ever used: medications patient was ever exposed to.
Meds at sampling: medications prescribed at time of inactive sample.

Medication abbreviations: PO, oral prednisone; IVMP, IV solumedrol; MTX, methotrexate; MMF, mycophenolate mofetil; HCQ, hydroxychloroquine; RTX, rituximab; CSA, cyclosporine; IVIG, IV immunoglobulin.

<u>Differentially expressed genes – ST3, ST6, ST12, and ST15.</u>
gene_id: Ensembl gene ID.
symbol: gene symbol.
gene_biotype: Ensembl gene biotype annotation.
FoldChange: DESeq2 calculated gene fold-change.
pval: DESeq2 p-value (Wald test).
qval: false-discovery rate corrected p-value.
baseMean: DESeq2 base mean value.
log2FoldChange: DESeq2 calculated log2 fold-change.
lfcSE: DESeq2 log2 fold-changer standard error.
stat: DESeq2 Wald test statistic.

<u>gProfileR pathway analysis – ST4, ST5, ST7, ST8, ST9, ST10, ST11, ST13, ST14, ST16, ST17, ST20, ST22, ST23, ST25, ST26, ST28, ST29.</u>
adjusted_p_value: g:SCS adjusted p-value
source: known pathway source.
term_id: Identifier number for the pathway.
term_name: name of the known pathway.
symbols: list of gene symbols in the category.
intersections: list of gene_ids in the category.

<u>List of genes increased in all 3 tissues – ST18.</u>
gene_id: Ensembl gene ID.
symbol: Ensembl symbol.
gene_biotype: Ensembl annotated biotype.

<u>WGCNA trait correlation analysis – ST19, ST21, ST24, ST27,</u>
gene_id: Ensembl gene ID.
symbol: Ensembl gene symbol.
gene_biotype: Ensembl annotated gene biotype.
moduleColor: WGCNA assigned module color.
GS.corr.Status: WGCNA biweight midcorrelation.
p.GS.Status: WGCNA p-value correlation.
direction: Directionality of the gene-gene correlation.

<u>Genes differentially expressed and trait-correlated – ST30</u>
gene_id: Ensembl gene ID.
symbol: Ensembl symbol.
gene_biotype: Ensembl annotated gene biotype.

<u>Gephi network statistics – ST31.</u>
gene_id: Ensembl gene ID.
symbol: Ensembl symbol.
gene_biotype: Ensembl gene biotype annotation.
trait: List of clinical traits the gene is associated with.
traitcount: Count of the number of clinical traits the gene is correlated with.
Degree – eigencentrality: Gephi network statistics.

SOMA untreated v control – ST32
Shows overlap between previously published SOMA results and the current study for untreated JDM vs. control sera.

gene_id: Ensembl gene ID.
symbol: Ensembl symbol.
gene_biotype: Ensembl gene biotype annotation.
Uniprot: Uniprot ID of the SOMA target
Effect direction: Compares the fold-change between PBMC transcriptomes and serum proteins.
PBMC RNA fold-change: The fold-change in PBMCs as untreated JDM / control.
PBMC RNA qval: DESeq2 adjusted p-value.
Plasma protein fold-change: The fold-change in sera by SOMA technology (untreated JDM / control).
Plasma protein qval: The adjusted p-value for the previous SOMA study.
SOMA result ID: row number of the original supplementary data from SOMA publication.

SOMA untreated v inactive – ST33
Shows the overlap between previously published SOMA results and the current study for untreated JDM vs. clinically inactive JDM.

gene_id: Ensembl gene ID.
symbol: Ensembl symbol.
gene_biotype: Ensembl gene biotype annotation.
PBMC RNA fold-change: transcriptome fold-change for untreated JDM / clinically inactive JDM.
PBMC RNA qval: adjusted p-value for the transcriptome fold-change.
PBMC RNA treatment response: Whether the PBMC RNA responded to treatment.
Uniprot: Uniprot ID for the SOMA target.
Plasma protein fold-change: Fold-change for sera protein by SOMA from previous publication for untreated JDM / clinically inactive JDM.
Plasma protein qval: adjusted p-value for the plasma fold-change.
Plasma protein treatment response: whether the plasma protein responded to treatment.
Response concordance: whether the PBMC RNA and sera protein agreed with respect to treatment response.

### 7. Supplementary Methods

#### 7.1. Gene-trait network analysis with WGCNA

The weighted-gene co-expression network analysis (**WGCNA**) was designed using the tutorials available from the WGCNA website. It is worth noting that WGCNA uses matrix math operations, and particularly for a full transcriptome with several samples, it was exceedingly slow using a standard R installation. We installed the Intel optimized math kernel library (MKL; 2018 version 2.199), and recompiled R to use this library with gcc version 7.2.0 using the following options:

```
1) source /opt/intel/mkl/bin/mklvars.sh intel64

2) MKL="-Wl,--no-as-needed -lmkl_gf_lp64 -Wl,--start-group -lmkl_gnu_thread  -lmkl_core  -Wl,--end-group -fopenmp  -ldl -lpthread -lm"

3) ./configure \
CFLAGS="-m64 -O3 -mtune=core2 -D_GLIBCXX_USE_CXX11_ABI=0" \
CXXFLAGS="-m64 -O3 -mtune=core2 -D_GLIBCXX_USE_CXX11_ABI=0" \
FFLAGS="-m64 -O3 -mtune=core2 -D_GLIBCXX_USE_CXX11_ABI=0" \
FCFLAGS="-m64 -O3 -mtune=core2 -D_GLIBCXX_USE_CXX11_ABI=0" \
--enable-R-shlib --with-blas="$MKL" --with-lapack
```

It also required manually soft-linking the shared-libraries to a system path first. However, the use of MKL libraries reduced the matrix math steps to a minute or two, a drastic improvement over the vanilla R version.

**7.2. Propagating standard deviation for RT-qPCR results**

For each sample tested, we used three separate technical replicates each of *otoferlin* target gene and beta actin control. We calculated the fold-change with the standard delta-delta Cq method (Cq = quantification cycle per MIQE guidelines), but appropriately combined the standard deviations at each step, e.g. the mean delta Cq was equal to the mean of the target minus the reference. The standard deviation of the delta Cq was the square root of the sum squared standard deviations of target and reference. This propagation of errors in ddCq is discussed at length in the supplement of a previous paper [1].

### 8. Supplementary Results

#### 8.1. Sex-linked differential expression in PBMCs

For unpaired DESeq2 analysis, we included sex as a covariate to avoid confounding due to sex imbalance in the available sample design. To show we were able to appropriately control for sex, we also calculated the reverse: accounted for disease status as a covariate and calculating for sex-linked differential expression. Only 35 genes were differentially expressed in this comparison. The top DE gene was *XIST* (X chromosome), which had -133-fold expression in males compared to females. Other top genes included *TTTY15* (343.4 FC; Y chromosome), *RPS4Y1* (262.9 FC; chromosome Y), *PRKY* (42.7 FC; chromosome Y), *ZFY* (60.8 FC; chromosome Y), *KDM5D* (293.5 FC; chromosome Y), and *TSIX* (-35.2 FC; chromosome X). We used these differentially expressed genes as input for principal components analysis in R with the prcomp function.

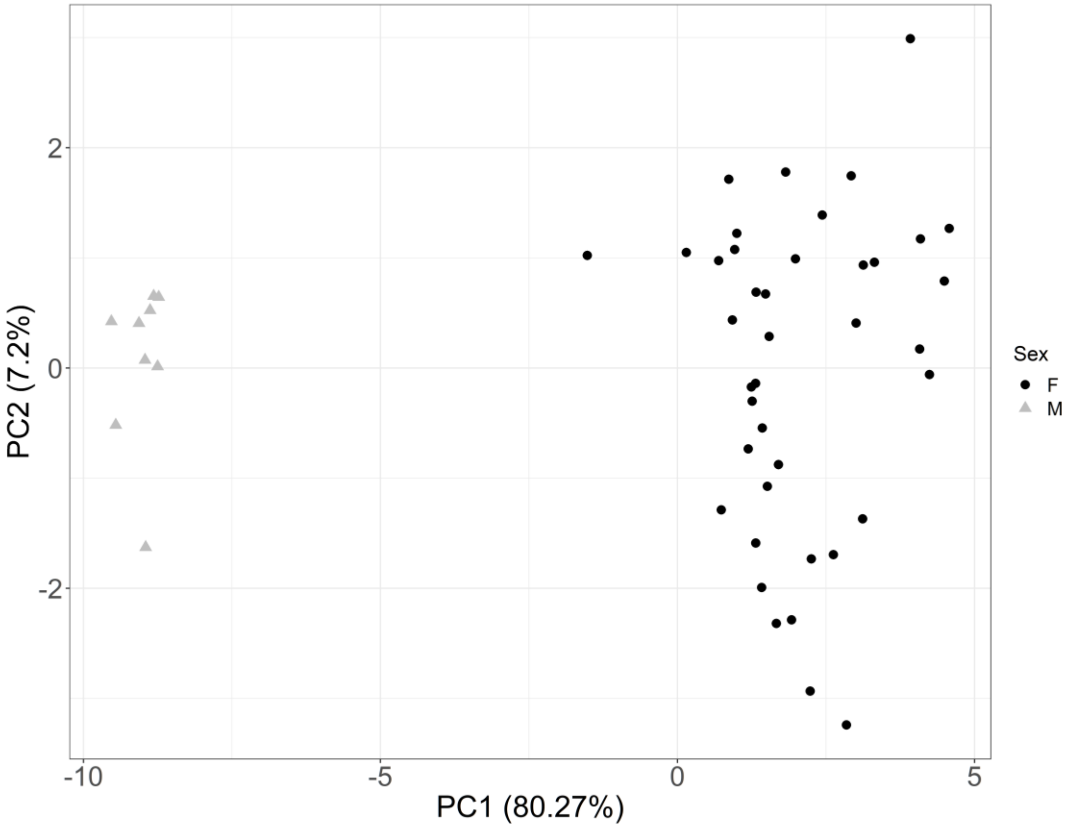


**Supp. Fig. 1 – Sex-linked genes perfectly separate XX and XY samples**
The top two PCs are on the x-axis and y-axis. Each dot is an individual sample. Samples expected to be XX are dark circles and samples expected to be XY are light grey triangles. The majority of variance (80.3%) is explained by the first PC, which perfectly separates samples by sex.

It is important to note that biological sex, genetic sex as determined by number / composition of sex chromosomes, and gender are not equivalent. These data show that the samples all had the expected genetic sex chromosomes based on the clinically reported anatomical sex.
